# Visual blur disrupts the kinematic and temporal aspects of reach-grasp-lift movements

**DOI:** 10.1101/2025.06.05.658186

**Authors:** William E. A. Sheppard, Carlo Campagnoli, Richard M. Wilkie, Rigmor C. Baraas, Rachel. O. Coats

**Author notes:** Corresponding Author: William Sheppard.

## Abstract

Degraded vision (caused by pathological reasons or monocular viewing) has been shown to affect fine motor control. However, there is a dearth of work examining the effects of “cataract-like” blur on reach-to-grasp performance. There is, however, a trend towards amblyopic blur being associated with deficits in reach-to-grasp performance, suggesting that timely intervention in treating cataracts is likely to be essential to maintain a functional ageing population. 18 participants performed a reach-to-grasp task. They reached for and precision grasped high and low-contrast cuboid targets under three visual conditions: binocular blur, monocular blur (full vision in the other eye) and full vision. They also performed contrast sensitivity, stereoacuity and visual acuity tests. Visual blur was associated with changes to the kinematics of prehensile movements’ early/acceleration stage (maximum acceleration and maximum velocity) and maximum grip aperture. Visual blur also caused the period from first contact with the target to the time it was lifted (dwell time) to be elongated. These results suggest that changes in prehension associated with visual blur are linked to differences in the planning and online control of prehension movements.

## Introduction

The ability to move one’s arm and hand towards an object to grasp it (prehension) is an essential component of many everyday actions, including reaching for a door handle or picking up a mug of tea. Therefore, factors affecting the execution of these movements can directly and significantly impact one’s quality of life. The spatiotemporal characteristics of prehensile movements depend on three fundamental aspects: the visual system’s internal state, the target object’s visual characteristics and the interaction between the two (1). To complete a prehensile movement, the individual must use their representation of these three aspects to estimate the hand and target positions (2) before planning the path between the two, as well as the final orientation of the grip (3). Once an individual has initiated the movement, they must effectively control it to its completion, which ideally requires visual and proprioceptive feedback.

Given the central role of vision in planning and executing prehensile movements, it is essential to understand how degraded vision affects reaching and grasping actions. There has been extensive research demonstrating the impact of monocular occlusion on both the planning and execution (online control) of prehensile movements (4–9) and a significant number of studies investigating prehension under monocular and/or binocular visual conditions, such as in the case of amblyopia (1,10–16), age-related macular degeneration (AMD (17–21)) and glaucoma (17,20,22–24).

Grant and Conway (2019) documented the effect of monocular occlusion on the spatial and temporal aspects of prehension. Specifically, monocular occlusion was associated with decreased maximum velocity (MA), increased maximum grip aperture (MGA) and longer movement time, primarily caused by an extended deceleration phase and contact-to-lift duration (dwell time). Interestingly, the effect of monocular vision on MGA and dwell time disappeared when visual feedback during the movement was removed, suggesting that monocular vision directly impacts the online control of the movement. In contrast, the aspects of the movement related to planning (time to MV and the time to MGA) were not affected by the removal of visual feedback.

Similar research has been conducted in anisometropic amblyopia patients, although the findings were more heterogeneous. There is consensus that amblyopia is associated with an increase in total movement time (10,11,14) and dwell time (10,11), as well as a trend towards a decrease in MV (11,14). However, as shown in Table 1, other variables, including maximum acceleration (MA), duration of the acceleration phase, MV, duration of the deceleration phase, and MGA, do not show a consistent pattern or are inconsistently reported (10,11,14).

**Table 1.** A summary of the effects of amblyopia on kinematic markers in prehensile tasks. The effects in the table are those associated with amblyopia vs. controls, i.e. “Increased” suggests that amblyopia was associated with an increase in a variable relative to controls.

|  | Buckley et al.<br>(2015) | Grant et al.<br>(2007) | Niechwiej-Szwedo et al.<br>(2011) |
| --- | --- | --- | --- |
| Movement time | Increased | Increased | Increased |
| Maximum velocity | No significant<br>effect | Decreased | Decreased |
| Maximum acceleration | Not reported | Not reported | Decreased |
| Duration of the acceleration<br>phase | No significant<br>effect | Not reported | Decreased |
| Maximum deceleration | Not reported | Not reported | No significant<br>effect |
| Duration of the deceleration<br>phase | Not reported | Not reported | No significant<br>effect |
| MGA | Decreased | No significant<br>effect | Not reported |
| Dwell time | Increased | Increased | Not reported |

Reduced contrast between stimuli and background also appears to magnify amblyopia’s effects on prehension. For example, amblyopes showed a larger increase, relative to controls, in planning time and grip aperture at contact with low contrast targets relative to the high contrast targets (1). The performance deficits demonstrated in those with amblyopia did not correlate with reduced binocularity or VA, but reduced CS was associated with increased target localisation errors (1). This is not surprising as amblyopia and visual blur have been demonstrated to degrade CS by increasing spatial sensitivity thresholds across the spectrum of spatial frequencies (25–27), with the effect being particularly strong at higher spatial frequencies associated with fine depth perception (28).

At the point of writing, there is little to no research on the effects of cataracts and cataract removal surgery on prehension. According to The Royal College of Ophthalmologists 2022 report, over 450,000 cataract removal procedures were carried out in England in the financial year 2019/20, an 11% increase from three years earlier. To complicate matters, the medical history of cataract patients is highly variable and heterogeneous, raising concerns about the generalizability of the findings.

Therefore, in the present study, we aimed to systematically assess the effects of cataract-like visual blur on grasping, which, at the time of writing, has not been directly assessed in previous research. We manipulated healthy participants’ vision by artificially blurring one or both eyes’ visual field, since cataracts can develop in either eye (monocularly) or both eyes (binocularly), whilst they performed a reach-to-grasp task to high and low contrast targets. We predicted that increased blur would be associated with changes in the movement’s spatial and temporal aspects, particularly in the scaling of MGA, the duration of overall MT, the duration of the deceleration phase, and the duration of dwell time. Additionally, the effects of degraded vision would likely be magnified when the participants reached for low-contrast targets. These findings have the potential to contribute to the growing body of evidence for the importance of timely cataract removal.

## Methods

### Participants

An opportunity sample of 19 participants completed the study between 24/10/2023 and 07/11/2023. One participant repeatedly failed to follow the task instructions, so the analysis did not include their data.

The ages of the final sample of 18 participants (11 female, 7 male) ranged from 20 to 31 (*mean* = 24.23 years, *sd* = 3.36 years) years old. All participants reported having normal or corrected-to-normal vision, and all participants reported being right-handed.

Participants were excluded if they were under 18, had persistent low vision that could not be corrected with glasses, had any disorder/condition that may affect their ability to grasp objects/impair their coordination, or were left-handed. Participants gave written consent for their participation in the study. Upon completing the study, the participants were entered into a prize draw to win a £30 Amazon voucher. The University of Leeds School of Psychology Ethics Committee granted ethical approval on 12/05/2023 (Ethics Reference Number: PSYC-899).

### Design

The present study employed a within-subjects experimental design whereby participants completed four tasks (reach-grasp-lift [RGL], contrast sensitivity [CS], visual acuity [VA] and stereoacuity) under three visual conditions (full vision, monocular blur and binocular blur). Participants completed the RGL, CS, and VA tasks in the first testing session in that order, with the order of the visual conditions being counterbalanced. Due to an oversight by the researchers, stereoacuity data were not collected during the initial testing session and were obtained at a later testing session.

Within each visual condition, the RGL task was varied across two levels of contrast (‘high’, a wooden target against a black background; ‘low’, a wooden target against a wooden background) and three distances (150 mm, 250 mm, 350 mm), giving rise to a 3 (visual condition) x 3 (distance) x 2 (contrast) within-subject design. The participants repeated each condition eight times, giving 144 trials split into three blocks of 48 trials by visual condition. The experiment lasted 60 to 90 minutes in total.

### Procedure

Upon arrival, participants were presented with an information sheet outlining what would be involved in the study and allowed to ask the researcher any questions. The participants read and signed a consent form confirming their eligibility and willingness to participate in the study. Finally, participants completed the eye dominance test. The researcher then fitted the participants with the first pair of glasses (clear lenses over both eyes, a clear lens over the dominant eye and a blurred one over the other, or blurred lenses over both eyes), depending on the visual condition performed first.

Each participant sat as close to the table as possible with their navel in line with the midline of the table. They started each trial gripping the start point: a round wooden dowel at the near edge of the table, between their thumb and index finger, making sure all markers were visible to the cameras. Upon a go-signal, the participants reached for the target as quickly and accurately as possible, gripping the target dowels with their thumb and index finger before lifting it. The participant then held the target suspended in the air. The researcher stopped the recording before the participant replaced the target object on the table, and the trial ended. The participants repeated this procedure until they completed all the trials for that visual condition and were then taken to a separate room to complete the CS and VA tasks. This procedure was repeated for the three visual conditions before the participant was debriefed and allowed to leave.

In the second testing session, the participants completed the stereoacuity task once for each visual condition, in the same order as they had completed the initial testing session. The participant was then debriefed and allowed to leave.

### Materials and stimuli

The table used for the experiment was 1200 mm wide x 800 mm deep x 730 mm high and was painted black. A board was mounted on the table that was 608 mm wide x 826 mm deep x 8 mm high. The board was slid in the Y-plane to change the contrast condition without the participant moving.

Runners positioned at 456 mm from the midline of the table limited the board’s movement. The left half of the board was plain wood, and the right half was painted black. These created the low and high contrast conditions, respectively (the target was wooden). The layout of each half of the board was identical so that the midline of each half of the board was populated with a round wooden dowel of 27 mm diameter on the near edge (the starting point for each trial) and three raised rectangular grooves (target locations) at 177 mm, 277 mm and 377 mm from the near edge of the table (150 mm, 250 mm and 350 mm from the far edge of the starting point). See Figure 1 A for a photo of the experimental setup and Figure 1 B for a schematic of each half of the board.

**Figure 1.**
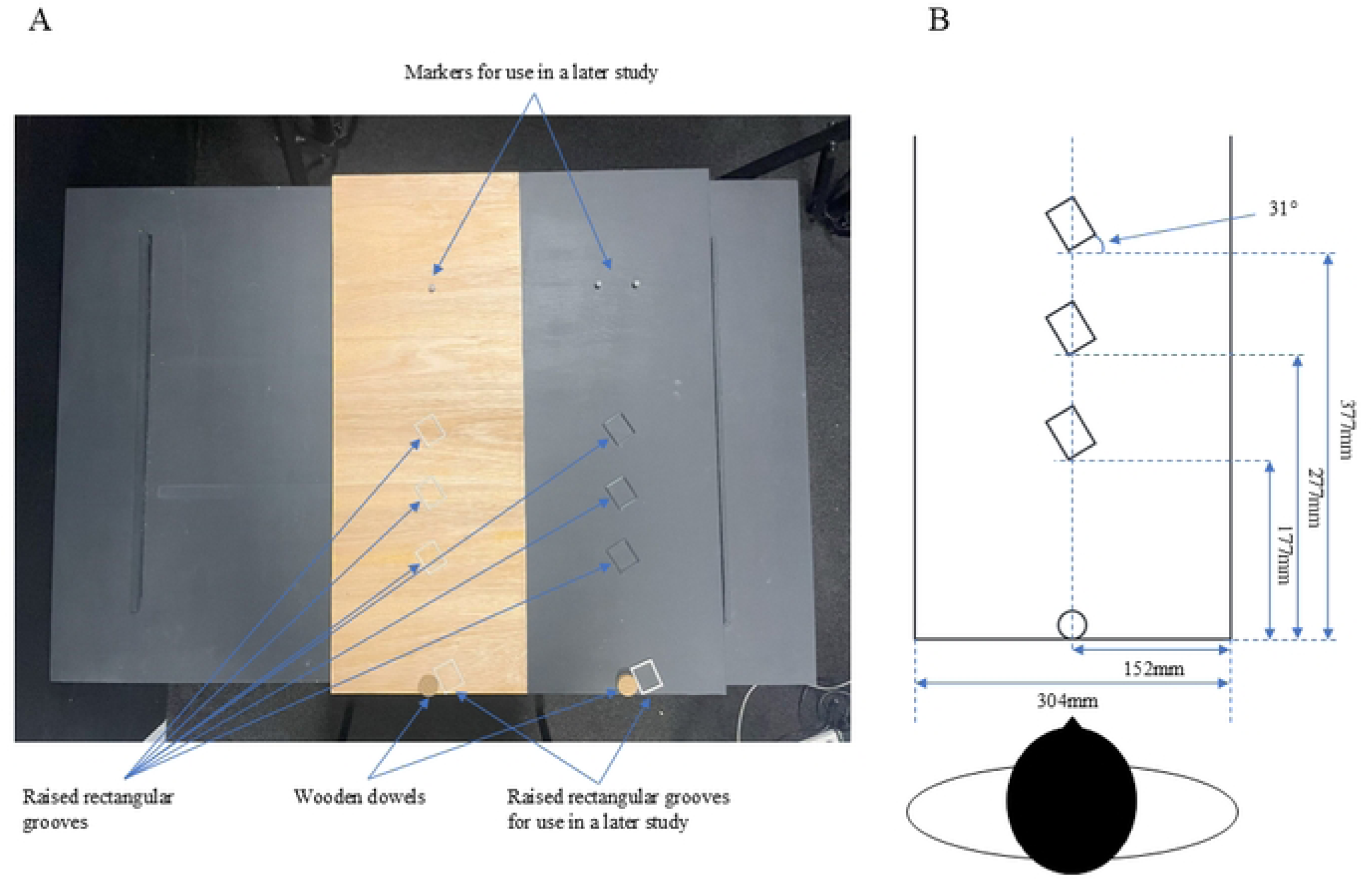
A, photo of the experimental setup used in the present study. B, a schematic of the experimental setup used in the present study.

Lightweight infrared reflective markers (12.7 mm diameters) were attached to the index fingernail, the index knuckle, the thumbnail and the wrist (head of the radius). The instantaneous position of the four markers and one placed on the top of the target were recorded at 120 Hz with submillimetre spatial resolution by four motion capture cameras (Optitrack Flex 13 [NaturalPoint, Corvallis, OR, USA]) The target objects were cuboids (29 mm wide x 36 mm depth x 86 mm height) with dowels fitted to two opposite sides (7 mm length x 10 mm diameter), as shown in Figure 2. These were positioned on the board so that the front edge subtended an angle of 31 degrees to the front of the board. During a pilot study, this angle was determined to give participants a comfortable reach.

**Figure 2.**
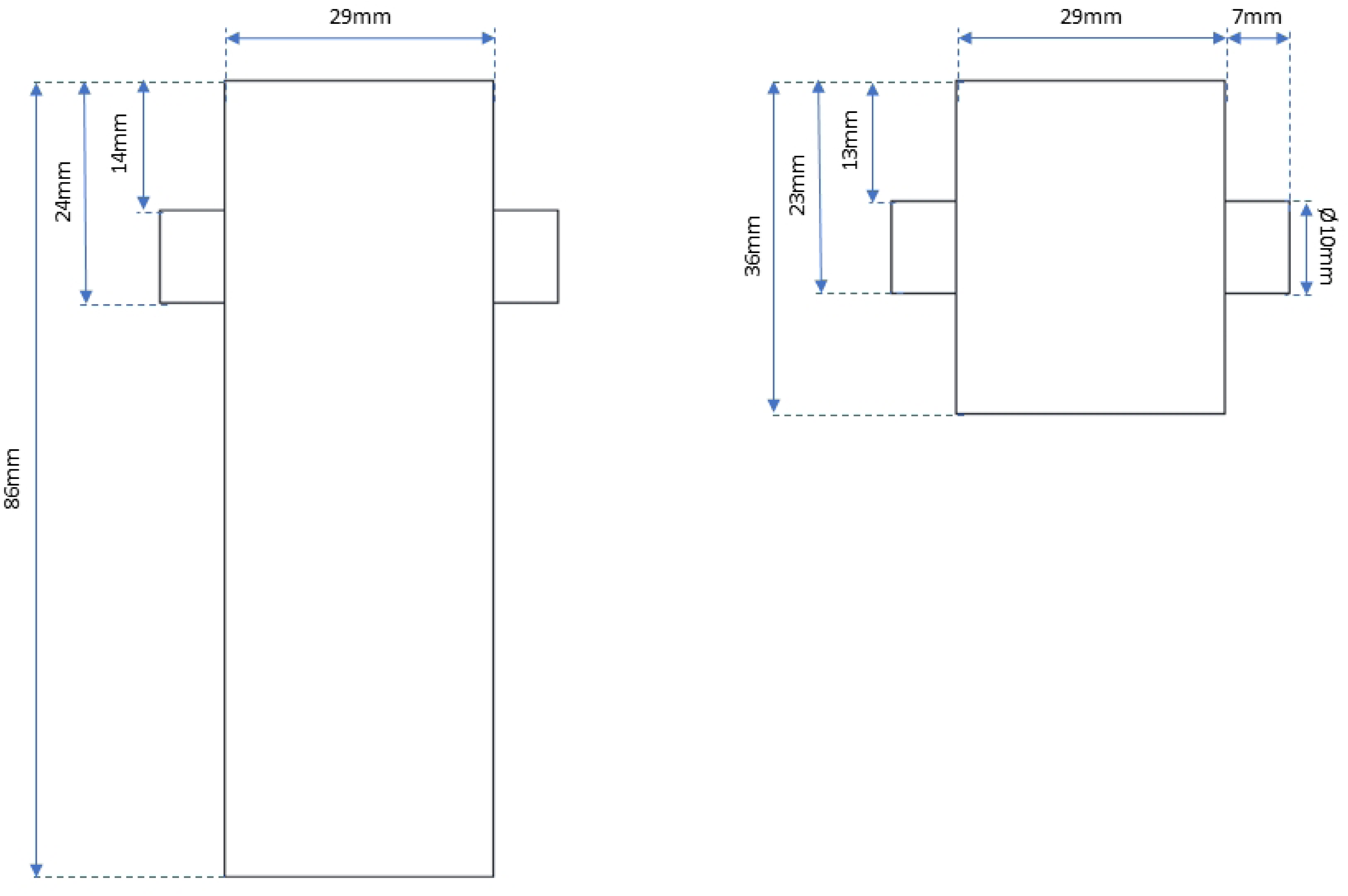
Target schematic from the front (left) and top (right) elevations.

Contrast sensitivity (CS) was measured using a right and left-eye Pelli-Robson CS chart (Pelli et al., 1988) calibrated at a 1m distance. Visual acuity (VA) was measured using the Bailey Lovie Chart 4, calibrated at a 6 m distance (30). Stereoacuity was measured at 40 cm using the TNO stereoacuity test (31,32). Eye dominance was determined using an ‘alignment test’ (33).

### Statistical analysis

The effect of the visual conditions on CS, stereoacuity, and VA was estimated using a standardised procedure after calculating the thresholds. A multilevel approach (using a Generalised Linear Mixed Model; GLMM) was taken (34). For each vision test score, the visual condition was entered into the model as a fixed factor, with a random intercept within each participant to model individual differences in each visual measure.

A similar GLMM approach was taken for the RGL data. Each outcome measure (listed and defined below in Table 2) was modelled as a function of three fixed factors: visual condition, contrast and distance. The maximal model included the three main effects plus all the two-way and the three-way interactions and covariates (the CS, VA, stereoacuity and session number). If a fixed effect was not significant and not included in the minimal model (the main effects of visual condition, contrast, distance, covariates and the two-way interaction of visual condition and contrast), it was removed from the model in order to achieve a parsimonious model; these are represented by a “–” symbol in the model tables. Some non-significant predictors excluded from the minimal model can still be seen in Tables 3, 4 and 5; this is due to each interaction effect containing several comparisons; for example, if we look at the two-way interaction of contrast and distance, this includes two comparisons: contrast x near vs. mid distance and contrast x near vs. far distance. If contrast x near vs. mid distance produces a significant effect and contrast x near vs. far distance does not, both comparisons will remain in the final model table, with the non-significant results represented as *NS*.

**Table 2.**
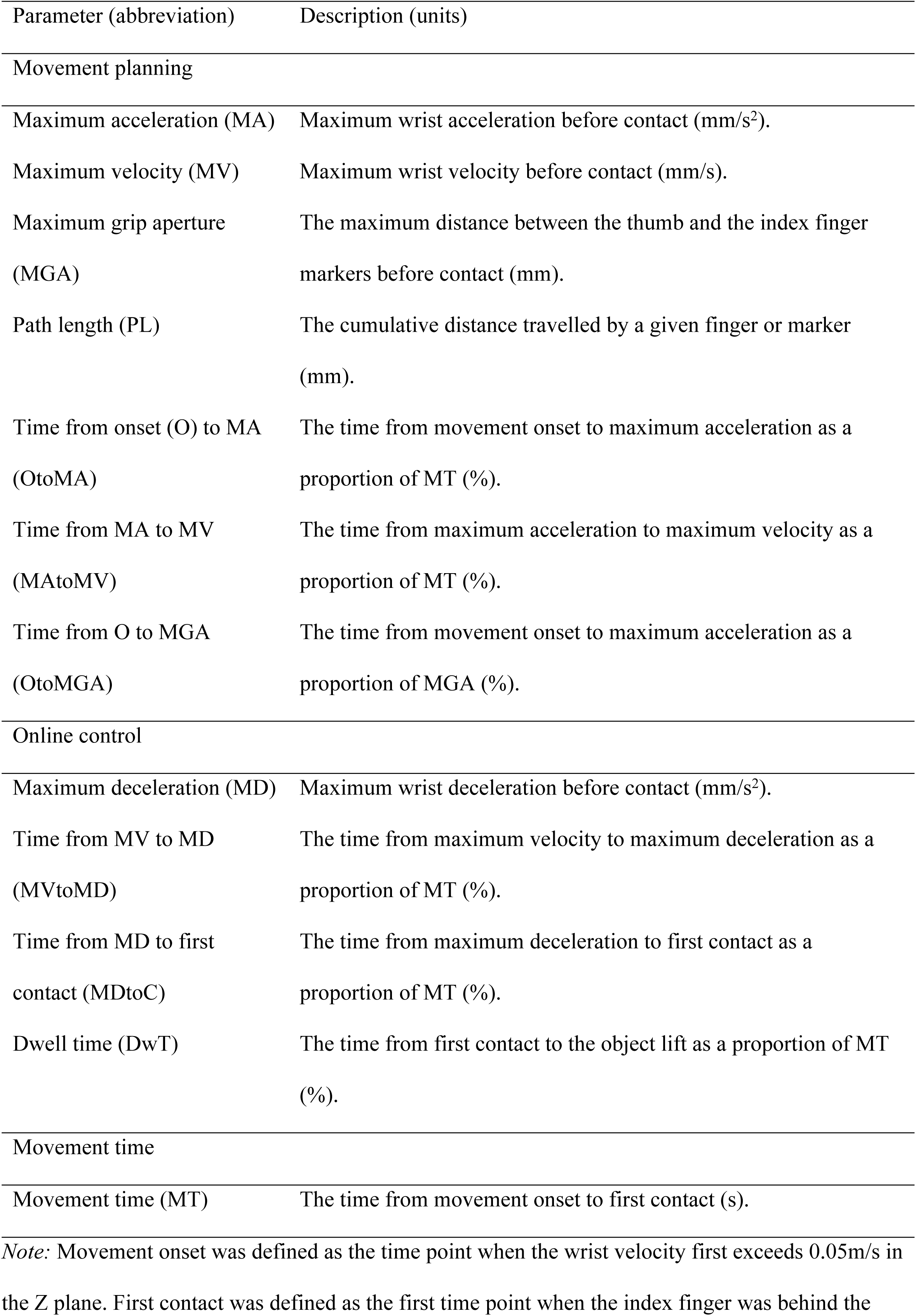
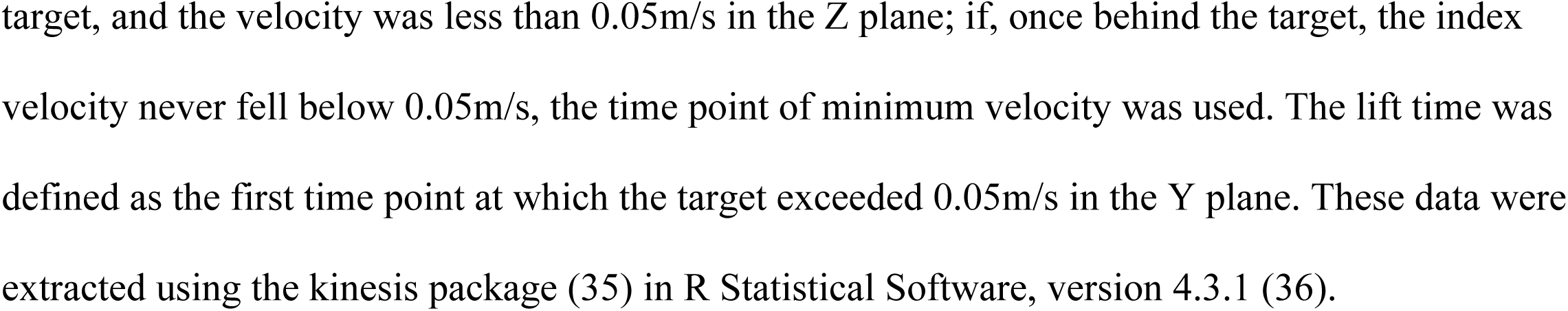
The definitions of the kinematic and timing measures and their abbreviations.

| Parameter (abbreviation) | Description (units) |
| --- | --- |
| Movement planning |  |
| Maximum acceleration (MA) | Maximum wrist acceleration before contact (mm/s<sup>2</sup>). |
| Maximum velocity (MV) | Maximum wrist velocity before contact (mm/s). |
| Maximum grip aperture (MGA) | The maximum distance between the thumb and the index finger markers before contact (mm). |
| Path length (PL) | The cumulative distance travelled by a given finger or marker (mm). |
| Time from onset (O) to MA (OtoMA) | The time from movement onset to maximum acceleration as a proportion of MT (%). |
| Time from MA to MV (MAtoMV) | The time from maximum acceleration to maximum velocity as a proportion of MT (%). |
| Time from O to MGA (OtoMGA) | The time from movement onset to maximum acceleration as a proportion of MGA (%). |
| Online control |  |
| Maximum deceleration (MD) | Maximum wrist deceleration before contact (mm/s<sup>2</sup>). |
| Time from MV to MD (MVtoMD) | The time from maximum velocity to maximum deceleration as a proportion of MT (%). |
| Time from MD to first contact (MDtoC) | The time from maximum deceleration to first contact as a proportion of MT (%). |
| Dwell time (DwT) | The time from first contact to the object lift as a proportion of MT (%). |
| Movement time |  |
| Movement time (MT) | The time from movement onset to first contact (s). |

*Note:* Movement onset was defined as the time point when the wrist velocity first exceeds 0.05m/s in the Z plane. First contact was defined as the first time point when the index finger was behind the
target, and the velocity was less than 0.05m/s in the Z plane; if, once behind the target, the index velocity never fell below 0.05m/s, the time point of minimum velocity was used. The lift time was defined as the first time point at which the target exceeded 0.05m/s in the Y plane. These data were extracted using the kinesis package (35) in R Statistical Software, version 4.3.1 (36).

**Table 3.** MLM estimates for fixed and random effects predicting performance on contrast sensitivity, stereoacuity and visual acuity tasks.

| Estimate | Contrast sensitivity | Stereoacuity | Visual acuity |
| --- | --- | --- | --- |
| Intercept | 1.80***<br>[1.77, 1.83] | 1.01***<br>[0.69, 1.34] | 1.07***<br>[0.94, 1.19] |
| Fixed effects ($\beta$) | | | |
| Binocular blur vs. monocular blur | 0.18***<br>[0.14, 0.21] | -0.57**<br>[-0.91, -0.24] | -0.28***<br>[-0.31, -0.25] |
| Monocular blur vs. full vision | 0.12***<br>[0.08, 0.15] | -0.39**<br>[-0.62, -0.16] | -0.18***<br>[-0.21, -0.15] |
| Random effects ($\sigma$) | | | |
| Participant ID | 0.04 | 0.43 | 0.12 |
| Model fit |  |  |  |
| R2 marginal | 0.668 | 0.077 | 0.366 |
| R2 conditional | 0.872 | 0.399 | 0.859 |

Note: The participant's performance on the stereoacuity tasks was standardised (i.e., with a mean equal to zero and *SD* equal to one). None of the outcome variables were normally distributed and were, therefore, modelled using the Gamma distribution. The Gamma distribution only accepts positive variables; thus, stereoacuity and visual acuity were made positive by adding 1. Due to the stereoacuity scores being centred, the $\beta$ values for the stereoacuity task are expressed as a proportion of the *grand SD* of stereoacuity (66.14 arcsecs). NS = $p \geq .05$ , \*\* = $p < .01$ , \*\*\* = $p < .001$ . The numbers in each cell represent the estimate [lower 95% CI, upper 95% CI].

**Table 4.**
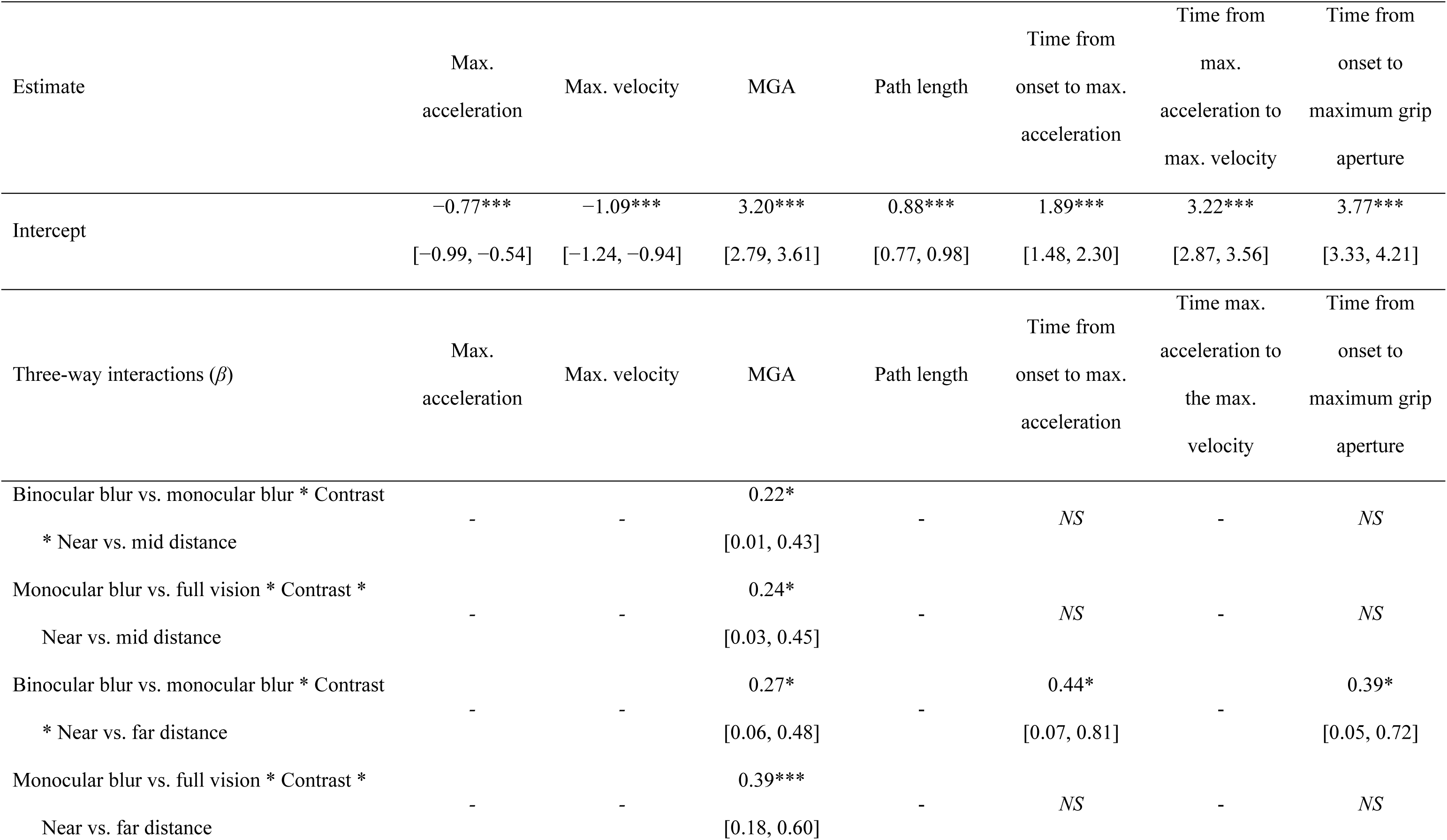
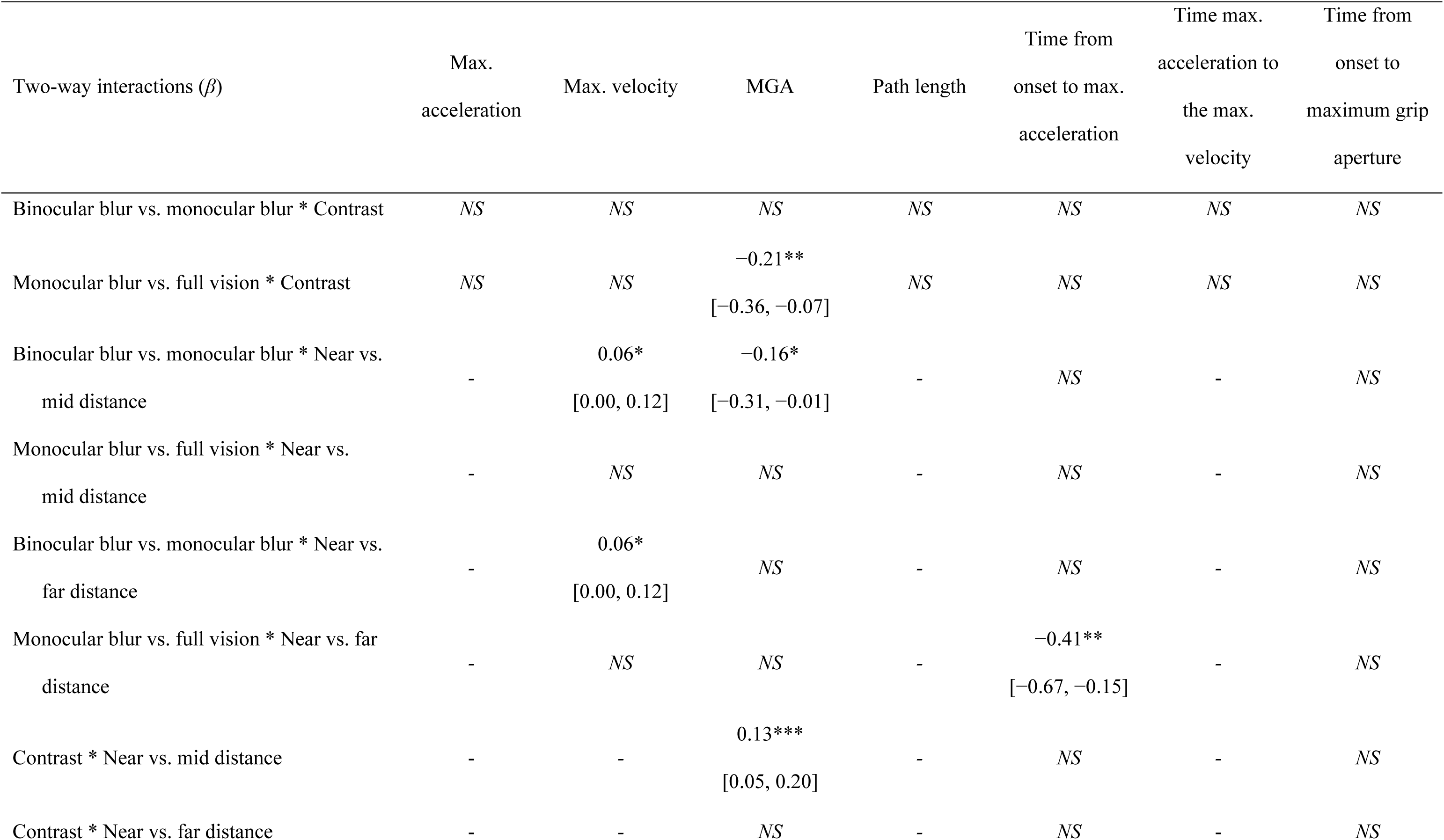
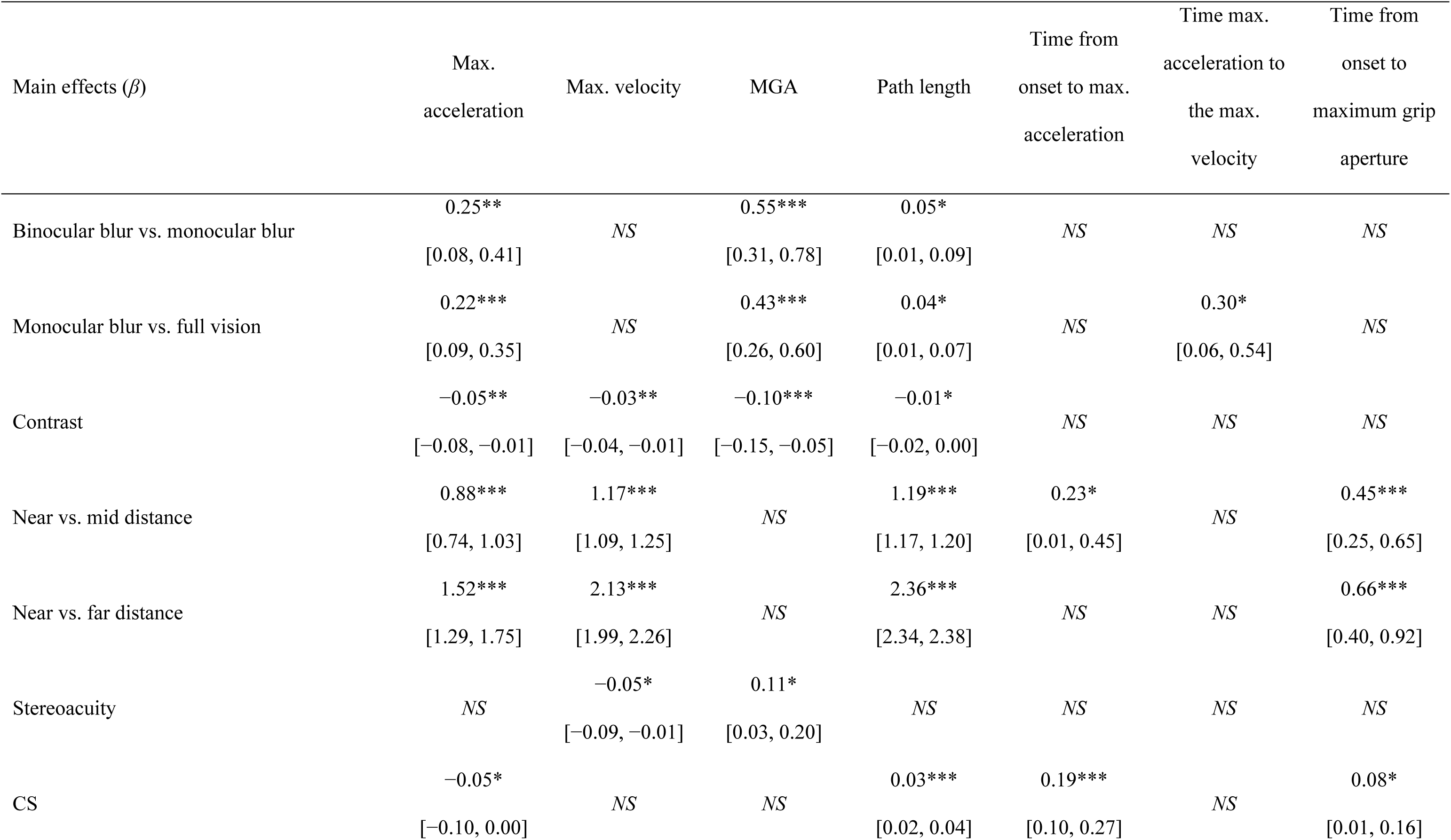
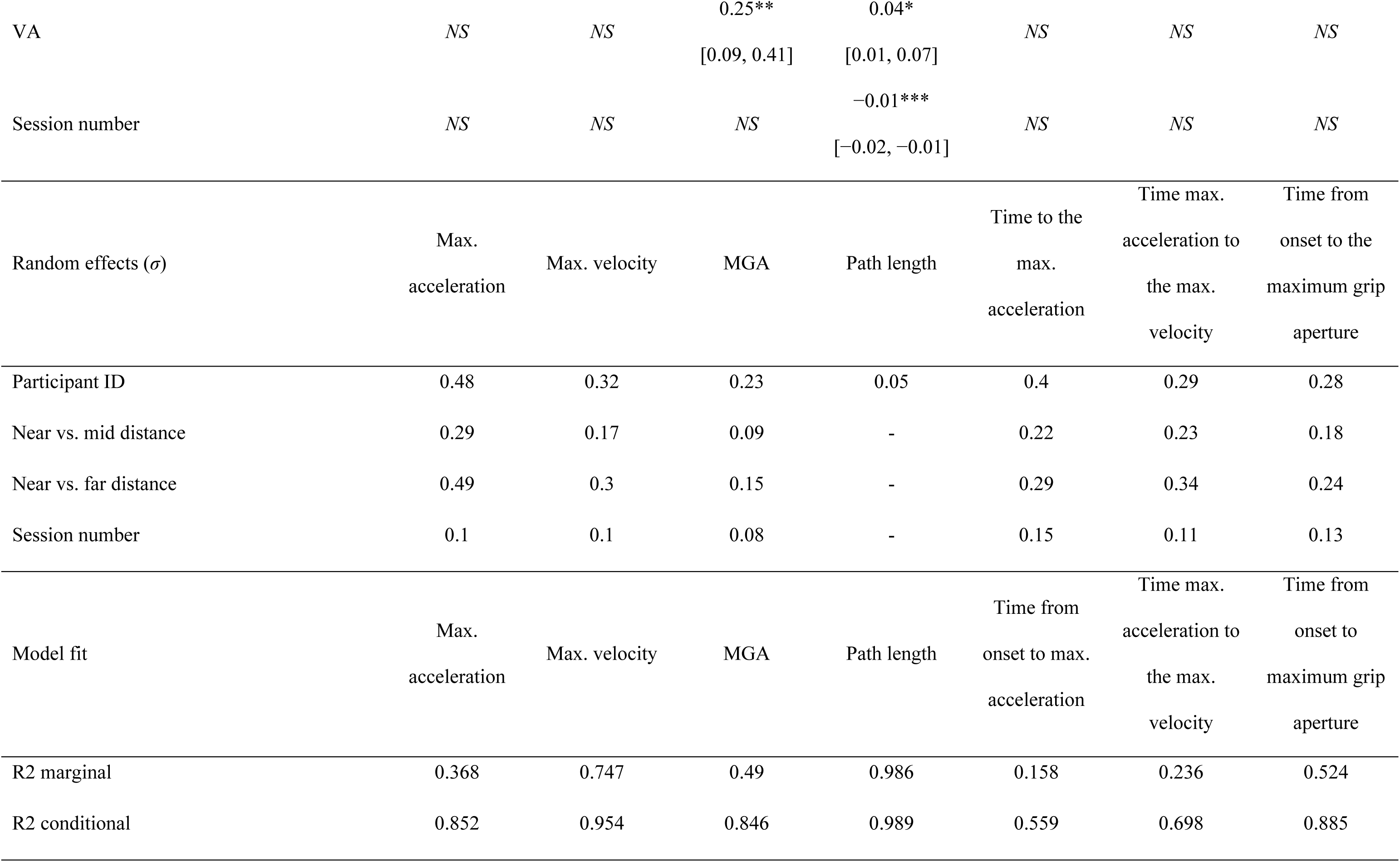
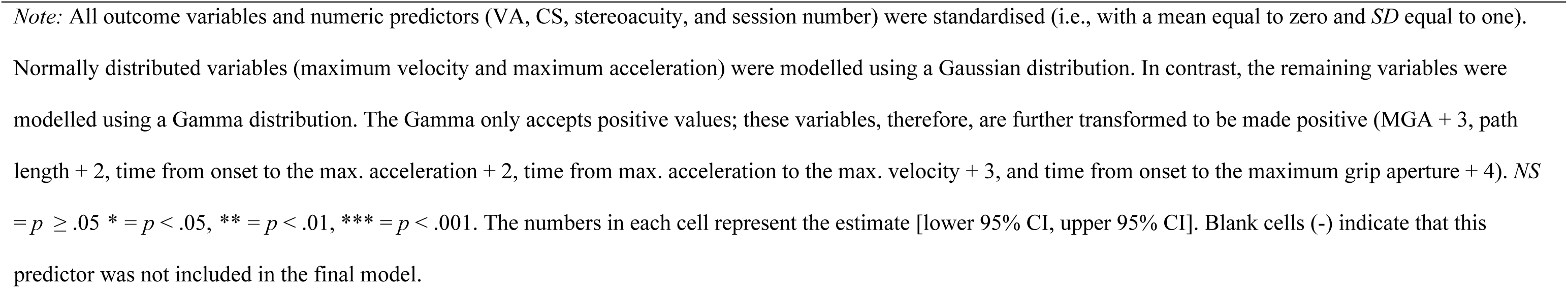
MLM estimates for fixed and random effects predicting movement planning of the RGL task.

**Table 5.** MLM estimates for fixed and random effects predicting online control characteristics of the RGL task.

| Estimate | Max.<br>declaration | Time from max.<br>velocity to the<br>max.<br>deceleration | Time from max.<br>deceleration to<br>contact | Time from<br>contact to lift<br>(dwell time) |
| --- | --- | --- | --- | --- |
| Intercept | 0.79***<br>[0.59, 1.00] | 3.29***<br>[2.90, 3.68] | 2.55***<br>[2.24, 2.86] | 2.21***<br>[1.76, 2.66] |
| Two-way interactions ($\beta$) | Max.<br>declaration | Time max.<br>velocity to the<br>max.<br>deceleration | Time from max.<br>deceleration to<br>contact | Time from<br>contact to lift |
| Binocular blur vs. monocular<br>blur * Contrast | *NS* | *NS* | *NS* | *NS* |
| Monocular blur vs. full vision *<br>Contrast | *NS* | *NS* | *NS* | *NS* |
| Binocular blur vs. monocular<br>blur * Near vs. mid distance | - | *NS* | *NS* | - |
| Monocular blur vs. full vision *<br>Near vs. mid distance | - | -0.33**<br>[-0.57, -0.10] | 0.29**<br>[0.08, 0.50] | - |
| Binocular blur vs. monocular<br>blur * Near vs. far distance | - | *NS* | *NS* | - |
| Monocular blur vs. full vision *<br>Near vs. far distance | - | *NS* | *NS* | - |
| Contrast * Near vs. mid distance | 0.11*<br>[0.02, 0.19] | *NS* | - | - |
| Contrast * Near vs. far distance | *NS* | -0.24**<br>[-0.40, -0.08] | - | - |

| Main effects ($\beta$) | Max.<br>declaration | Time max.<br>velocity to the<br>max.<br>deceleration | Time from max.<br>deceleration to<br>contact | Time from<br>contact to lift |
| --- | --- | --- | --- | --- |
| Binocular blur vs. monocular<br>blur | *NS* | *NS* | *NS* | -0.53***<br>[-0.78, -0.29] |
| Monocular blur vs. full vision | *NS* | *NS* | *NS* | -0.39***<br>[-0.57, -0.20] |
| Contrast | *NS* | *NS* | *NS* | *NS* |
| Near vs. mid distance | -0.99***<br>[-1.15, -0.83] | *NS* | 0.51***<br>[0.28, 0.74] | -0.15 [-0.21,<br>-0.09]*** |
| Near vs. far distance | -1.50***<br>[-1.73, -1.27] | -0.38**<br>[-0.67, -0.10] | 1.01***<br>[0.68, 1.33] | -0.26***<br>[-0.32, -0.20] |
| Stereoacuity | -0.09*<br>[-0.17, -0.01] | -0.15*<br>[-0.29, 0.00] | *NS* | *NS* |
| CS | *NS* | *NS* | *NS* | *NS* |
| VA | *NS* | *NS* | *NS* | -0.19**<br>[-0.33, -0.05] |
| Session number | *NS* | *NS* | 0.08***<br>[0.05, 0.12] | -0.11***<br>[-0.14, -0.09] |
| Random effects ($\sigma$) | Max.<br>deceleration | Time max.<br>velocity to the<br>max.<br>deceleration | Time max.<br>deceleration to<br>contact | Time from<br>contact to lift |
| Participant ID | 0.43 | 0.32 | 0.25 | 0.37 |
| Near vs. mid distance | 0.32 | 0.22 | 0.22 | - |
| Near vs. far distance | 0.48 | 0.26 | 0.31 | - |
| Session number | 0.12 | 0.08 | - | - |

| Model fit | Max.<br>declaration | Time max.<br>velocity to the<br>max.<br>deceleration | Time from max.<br>deceleration to<br>contact | Time from<br>contact to lift |
| --- | --- | --- | --- | --- |
| R2 marginal | 0.381 | 0.406 | 0.559 | 0.203 |
| R2 conditional | 0.817 | 0.712 | 0.787 | 0.614 |

*Note:* All outcome variables and numeric predictors (VA, CS, stereoacuity, and session number) were standardised (i.e., with a mean equal to zero and *SD* equal to one). Maximum deceleration was normally distributed and was modelled using a Gaussian distribution. In contrast, the remaining variables were modelled using a Gamma distribution. The Gamma distribution only accepts positive values; therefore, these variables are further transformed to be positive (time from maximum velocity to maximum deceleration + 3, time from maximum deceleration to contact + 3 and time from contact to lift +2). None of the final models for the variables presented in this table contained a significant three-way interaction, so three-way interactions are not included. The numbers in each cell represent the estimate [lower 95% CI, upper 95% CI]. Blank cells (-) indicate that this predictor was not included in the final model. *NS* = $p \geq .05$ , \* = $p < .05$ , \*\* = $p < .01$ , \*\*\* = $p < .001$ .

The dependent variables were grouped into three categories to systematically examine how visual blur influences reach-to-grasp movements: movement planning, online control, and movement time. This classification reflects distinct phases of the prehensile movement, each associated with different aspects of motor execution and sensorimotor feedback. Movement planning variables primarily capture feedforward control mechanisms based on visual input before movement execution. They are labelled as follows: maximum acceleration [MA], maximum velocity [MV], maximum grip aperture [MGA], path length [PL], time from onset [O] to MA [OtoMA], time from MA to MV [MAtoMV], and time from O to MGA [OtoMGA]. Online control variables reflect adjustments made during movement execution. They are labelled as follows: maximum deceleration [MD], time from MV to MD [MVtoMD], time from MD to first contact [MDtoC], and dwell time [DwT]. Movement Time [MT] represents the overall temporal duration of the action and integrates aspects of movement planning and online control.

A random intercept was estimated for each participant to account for differences in vision and coordination between participants, and random slopes for distance and session number to account for variability in participant arm length and learning rates, respectively.

Regarding the time series analysis, six kinematic landmarks were selected to divide each reaching action: maximum acceleration (MA), maximum velocity (MV), maximum grip aperture (MGA), maximum deceleration (MD), first contact with the target (Contact) and when the participant lifts the target (Lift). These landmarks are shown in Figure 3 on an exemplar velocity curve.

**Figure 3.**
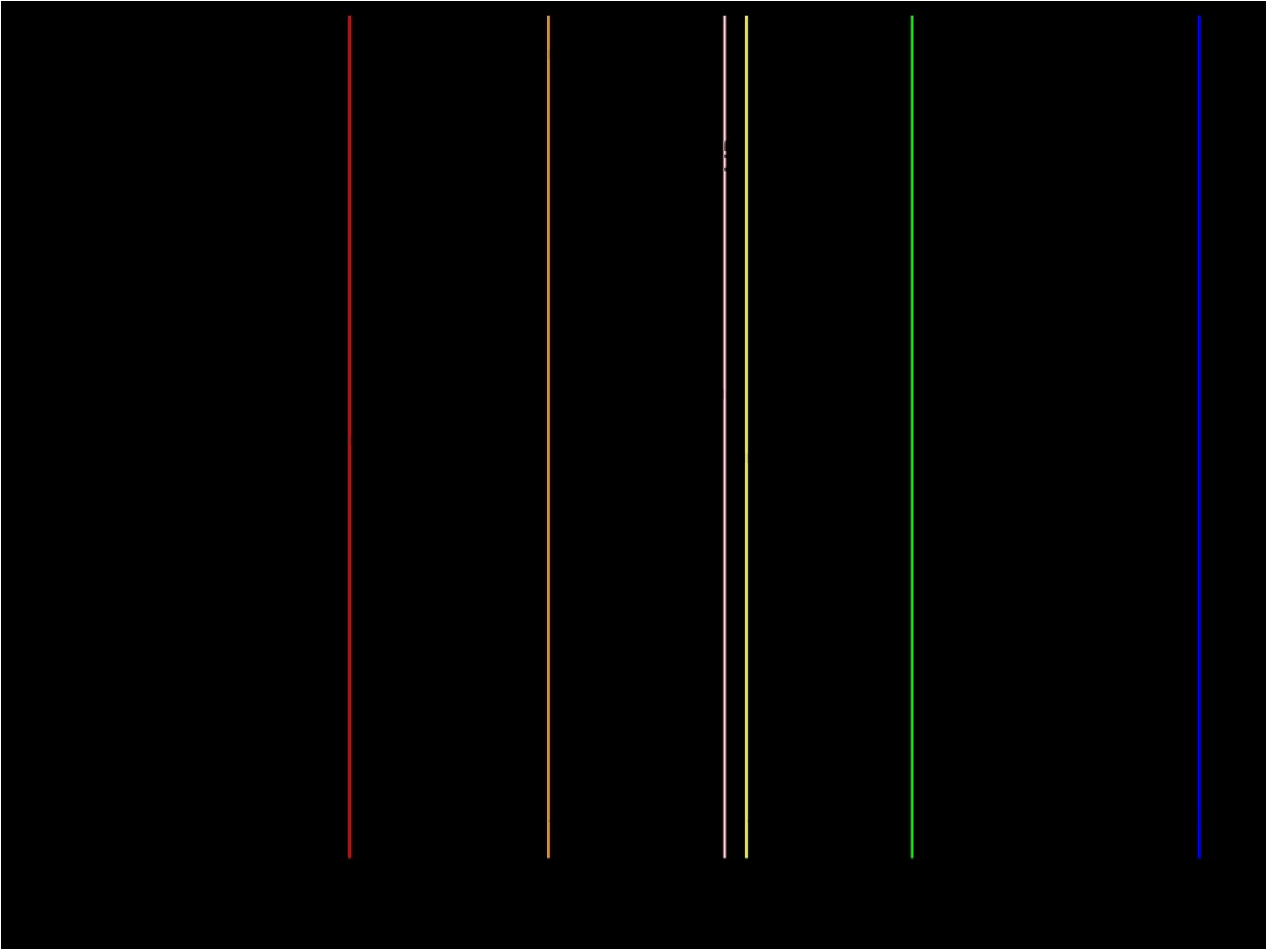
An example velocity profile showing the landmarks used to break down each movement.

The normally distributed outcome variables with no time component were modelled using a Gaussian distribution (MA, MV, and MD). In contrast, the remaining variables were modelled using the Gamma distribution because they have a zero-bound time component and are typically positively skewed, which aligns with the distribution’s assumptions. These variables were initially also modelled using the Inverse Gaussian distribution. However, after the performance of each model was assessed by comparing the BIC (Bayesian Information Criterion), all models were fitted using the Gamma distribution. A BIC difference greater than 10 gives “very strong” evidence that the model with the lower BIC value best fits the data, favouring models with lower numbers of factors (37,38). As all outcome variables were scaled to improve model fit, a small constant was added to each value (modelled with the Gamma or Inverse-Gaussian distributions), which only shifts the intercept and does not affect regression slopes (i.e. *β* coefficients) or significance tests. These analysis scripts are stored in a GitHub repository [https://github.com/willsheppard9895/blurNprehension], and the data is stored in the figshare repository [https://figshare.com/articles/dataset/Prehension_Data/28877057].

The outcome variables and numeric predictors (CS, stereoacuity, VA and session number) were centred and normalised. As the Inverse Gaussian and Gamma distributions only accept positive values, the outcome measures modelled using these distributions were further transformed to be positive (by adding the lowest integer to create a set of positive values; this was calculated for each variable), as per equation 1.

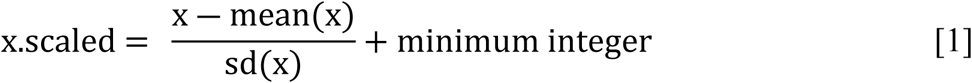

Visual condition, contrast, and distance were entered into each model as categorical predictors. Regarding visual condition, the participant’s performance was compared between binocular and monocular blur and between monocular blur and full vision to match the vision changes associated with first and second eye cataract surgery, respectively. Regarding distance, comparisons were made between near and mid distance and near and far distance. Significant three-way interactions are reported and plotted in the results section. Post-hoc tests were not conducted for significant three-way interactions due to the large number of pairwise comparisons (*N* = 153), which would greatly increase the likelihood of type I errors. Regarding significant two-way interactions, only those including visual condition will be reported and plotted in the results section. Any significant visual condition x contrast interactions will also be subject to post-hoc testing, as these interactions addressed the main aims of the study. Significant main effects of visual condition and contrast are also reported in the text. All MLM results are reported in Tables 3 (vision tests), 4 (movement planning), 5 (online control) and 6 (movement time). If one or more significant three-way interactions predict an outcome variable, the results section will not discuss any significant two-way interactions.

Inter-individual variation in each outcome measure was also assessed. For each outcome measure, the heterogeneity within the sample was assessed by comparing the relative size of the *SD* of the random intercepts allocated to each participant to the fixed intercept. As many of the intercepts were altered by the transformations applied to make them positive, this transformation was removed for this analysis. In the present case, this took the form β, where σ is equal to the magnitude of the random intercept *SD*, and *β* is equal to the magnitude of the fixed intercept minus the transformation. When this value exceeds 0.25, we concluded that the data are heterogeneous, as a participant at the 2.5th percentile would have a score equivalent to 0.5 of the mean, and a participant at the 97.5th percentile would have a score 1.5 times the mean (39). These results were only reported if the effect was heterogeneous. The random effects of session number and distance were not subject to this analysis, as these were entered to account for inter-individual variability in learning rate and arm length, respectively, and did not speak to the overall aims of the manuscript.

All analyses were performed using R Statistical Software (36). Kinematic measures were extracted from the RGB data using the Kinesis package (35), GLMMs were estimated using the lme4 package (40), p-values were estimated using Satterthwaite’s approximation through the lmerTest package (41), and estimated marginal means (*EMM*) for post-hoc testing were estimated using the emmeans package (42).

## Results

### Vision tests

To ensure that blurred conditions were affecting vision in a way similar to cataracts, an MLM analysis was carried out on the effects of visual conditions on contrast sensitivity (CS), stereoacuity, and visual acuity (VA) tasks. The *β* values in Table 3 predict the mean CS, stereoacuity and VA.

MLM analysis found significant effects of visual condition on CS (see Figure 4A), stereoacuity (see Figure 4B) and VA (see Figure 4C); monocular blur (relative to binocular blur) and full vision (relative to monocular blur) improved CS by 0.18 and 0.12 log units, stereoacuity by 37.93 and 25.79 arc secs and VA by 0.28 and 0.18 logMAR.

**Figure 4.**
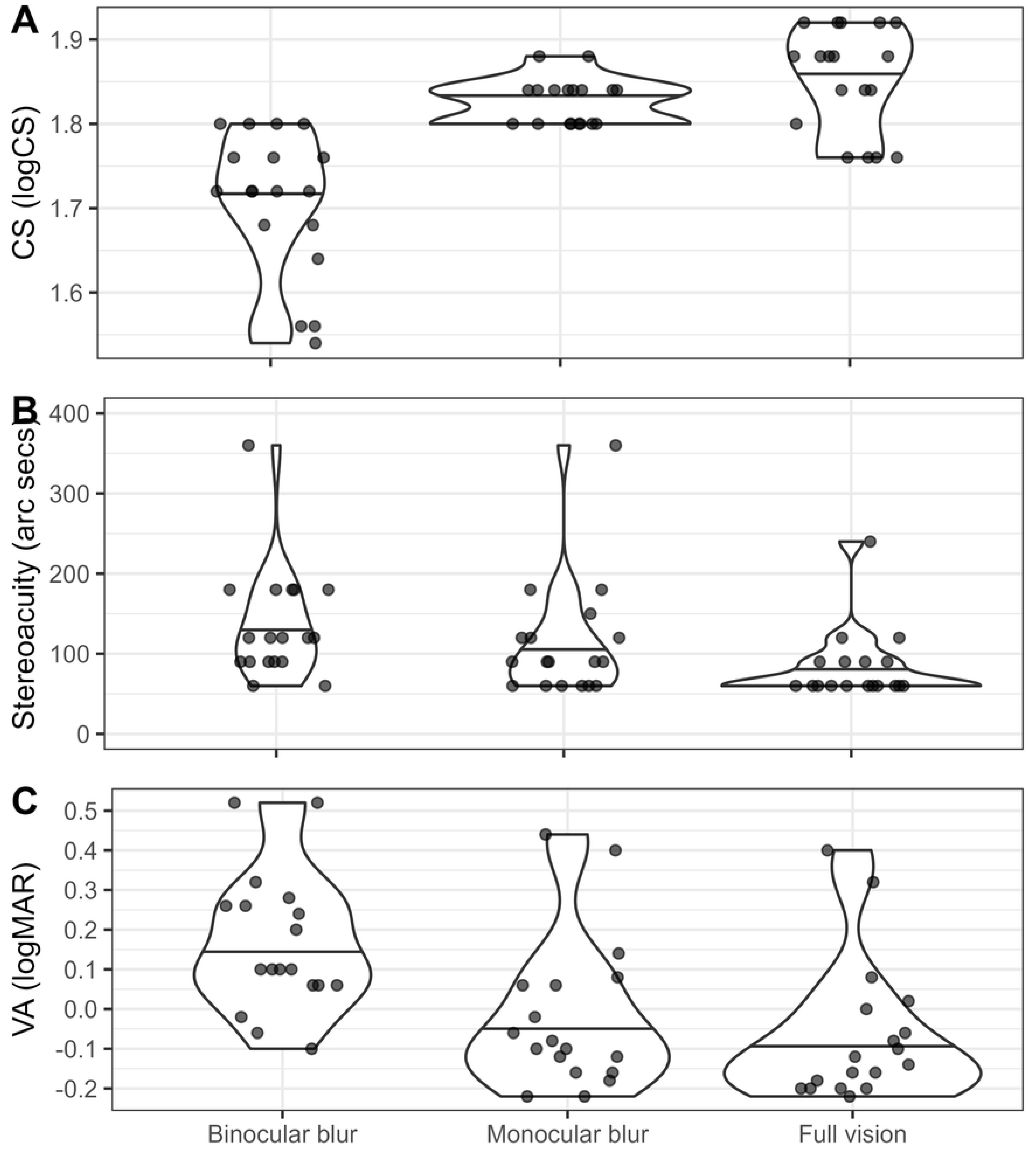
Effects of visual conditions on each vision test. The central black line of each violin represents the mean, and the outer lines represent the range. A. CS (log units). B. Stereoacuity (arc secs). C. VA (logMAR).

The *SD* of the random intercepts was 4300.00% and 171.43% of the fixed intercepts for stereoacuity and VA, respectively. This indicates extremely high individual differences in visual functions and that group-level averages may obscure meaningful variability influencing prehensile performance.

### Movement planning

The *β* values in Table 4 predict maximum acceleration, *µ_ma_*, maximum velocity, *µ_mv_*, maximum grip aperture (MGA), *µ_mga,_* path length, *µ_pl_*, the time from onset to maximum acceleration, *µ_2MA_*, the time from maximum acceleration to maximum velocity, *µ_MA2MV_*, and the time from onset to MGA, *µ_2MGA_*. As the outcome variables are centred, the *β* values are expressed as a proportion of the measure’s overall SD, e.g. *µ_ma_* is expressed as a proportion of the overall *SD* of MA (*SD_ma_*).

### Maximum acceleration

MLM analysis revealed significant main effects of visual condition and target contrast on maximum acceleration. Maximum acceleration increased progressively from binocular to monocular blur (67.09mm/s^2^ higher) to full vision (59.34 mm/s^2^ higher) and by 12.54 mm/s^2^ with high contrast compared to low contrast targets. The *SD* of the random intercept was 62.34% of the fixed intercept, reflecting moderate variability in baseline MA across participants.

### Maximum velocity

There were two significant two-way interactions between visual condition (binocular blur vs. monocular blur) x distance (near vs. mid) and visual condition (binocular blur vs. monocular blur) x distance (near vs. far). The increase in maximum velocity associated with monocular blur relative to binocular blur was greater at mid and far distances than near distances (see Figure 5). A significant main effect of contrast revealed that maximum velocity increased by 6.85mm/s with high contrast compared to low contrast targets.

**Figure 5.**
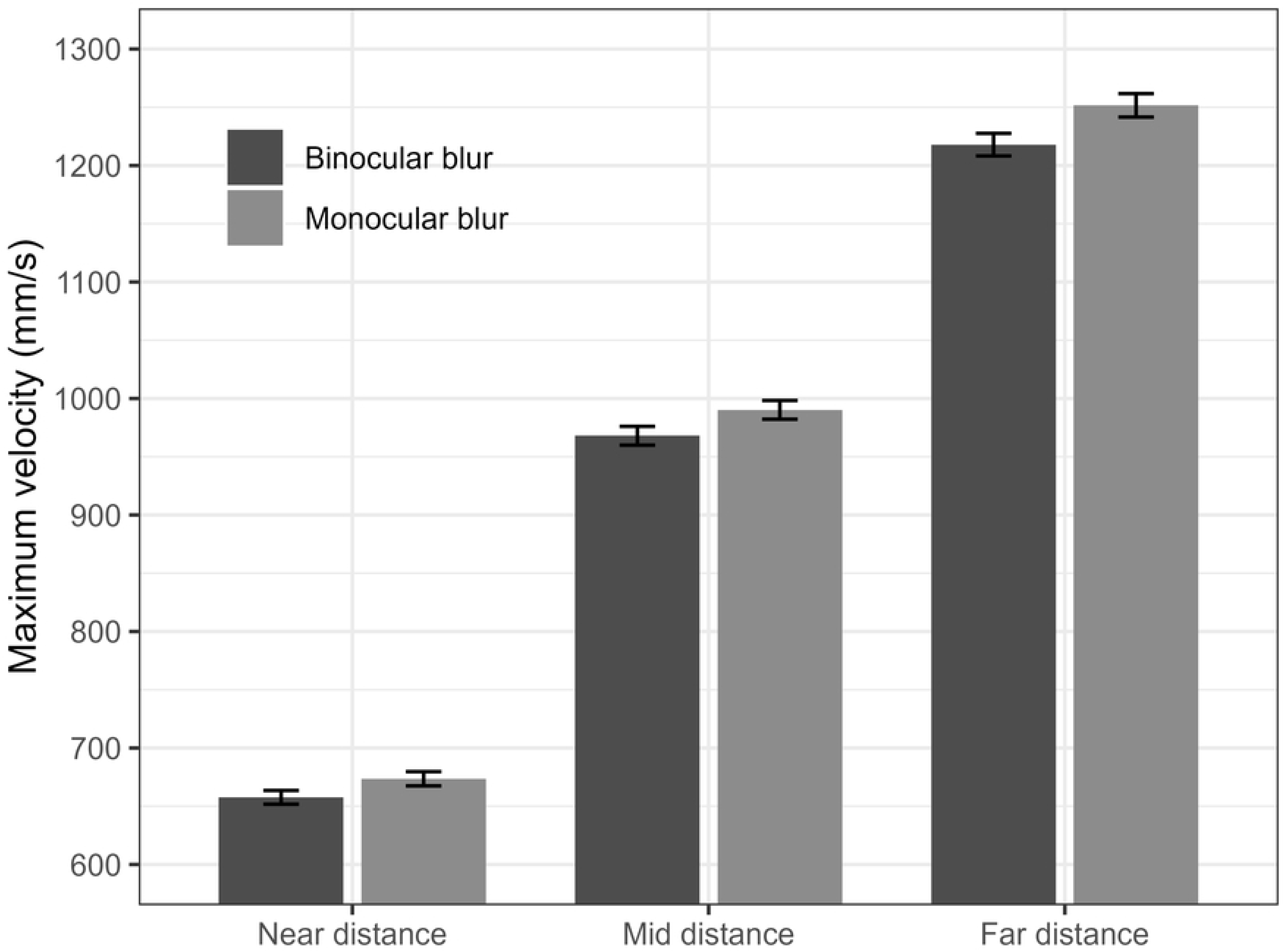
The effect of distance and visual condition (monocular and binocular blur) on maximum velocity (m/s). Error bars represent *standard error* (*SE*).

Participants’ baseline levels of MV showed variability, with the *SD* of the random intercepts estimated as 29.36% of the fixed intercept.

### Maximum Grip Aperture (MGA)

MLM analysis revealed four significant three-way interactions. When considering the interactions of visual condition (binocular blur vs. monocular blur) x contrast x distance, at both mid and far distances, monocular blur (relative to binocular blur) is associated with a decrease in MGA regardless of target contrast. However, at a near distance, monocular blur (relative to binocular blur) is associated with a reduction in MGA with low contrast targets and a slight increase or no difference in MGA with high contrast targets (Figure 6). Furthermore, when considering the interactions of visual condition (monocular blur vs. full vision) x contrast x distance, at both mid and far distances, full vision (relative to monocular blur) is associated with an increase in MGA regardless of target contrast. However, at near distance, full vision (relative to monocular blur) is associated with increased MGA with high contrast targets and a slight increase or no difference in MGA with low contrast targets (Figure 6).

**Figure 6.**
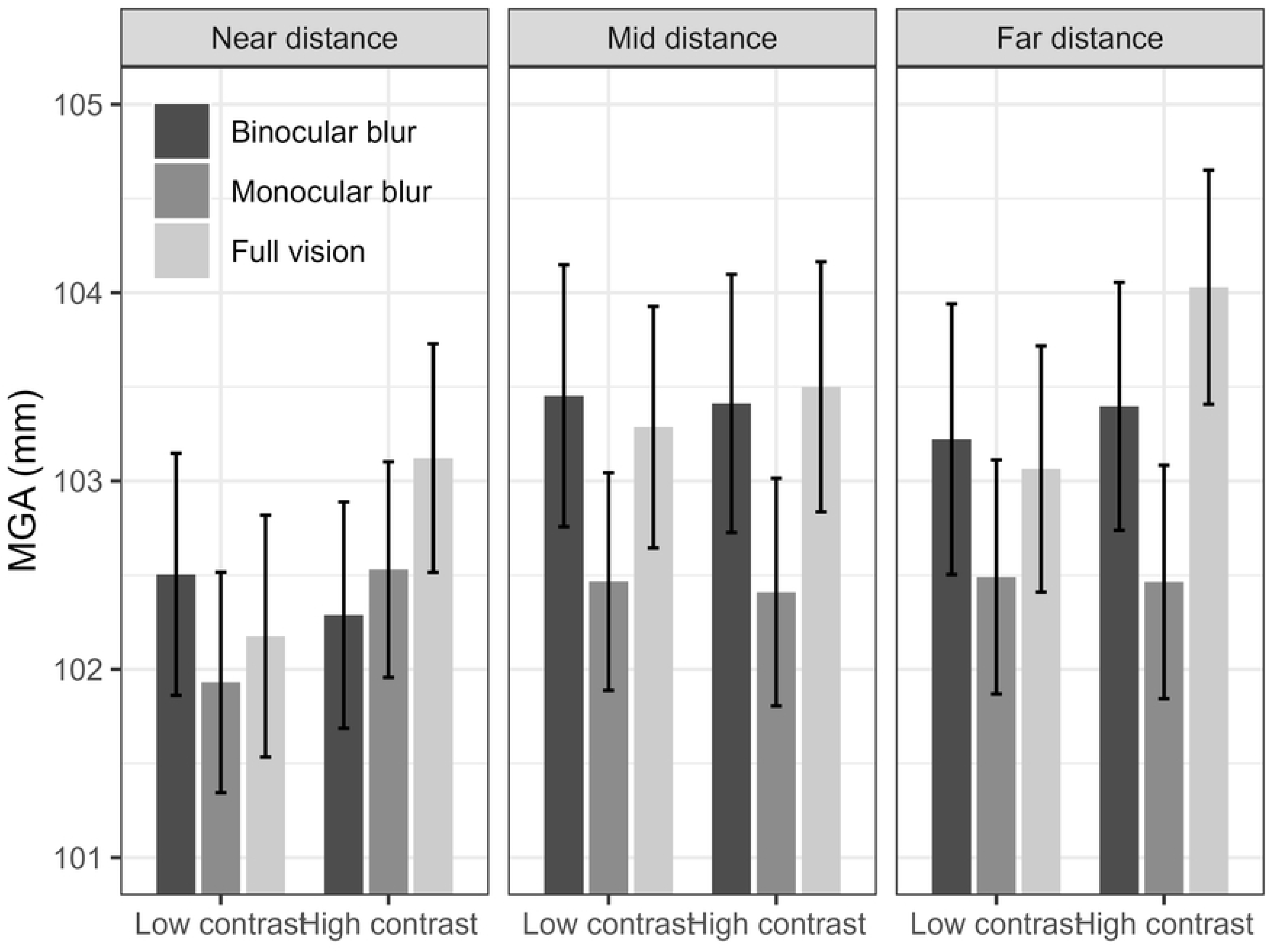
The effect of visual condition (binocular blur vs. monocular blur and monocular blur vs. full vision), contrast and distance (near vs. mid and near vs. far) on MGA (mm). Error bars represent *SE*.

MLM analysis revealed significant main effects of visual condition and contrast on MGA. MGA increased progressively from binocular to monocular blur (4 mm larger) to full vision (3.16 mm larger) and by 0.74 mm with high contrast compared to low contrast targets. The *SD* of the random intercepts was 115.00% of the fixed intercept, indicating high individual variability in base levels of MGA.

### Path length

MLM analysis revealed two significant main effects of visual condition on path length. Path length increased progressively from binocular to monocular blur (4.52 mm longer) to full vision (3.65 mm longer) and by 1.07 mm with high contrast compared to low contrast targets.

### Time to Maximum Acceleration (OtoMA)

A significant three-way visual condition (binocular blur vs. monocular blur) x contrast x distance (near vs. far) interaction indicates that at near distance, with low contrast targets, monocular blur is associated with a slight decrease in OtoMA; however, at far distance with low contrast targets, monocular blur is associated with a slight increase in OtoMA. Monocular blur is not associated with a change in OtoMA with high contrast targets at either distance (see Figure 7).

**Figure 7.**
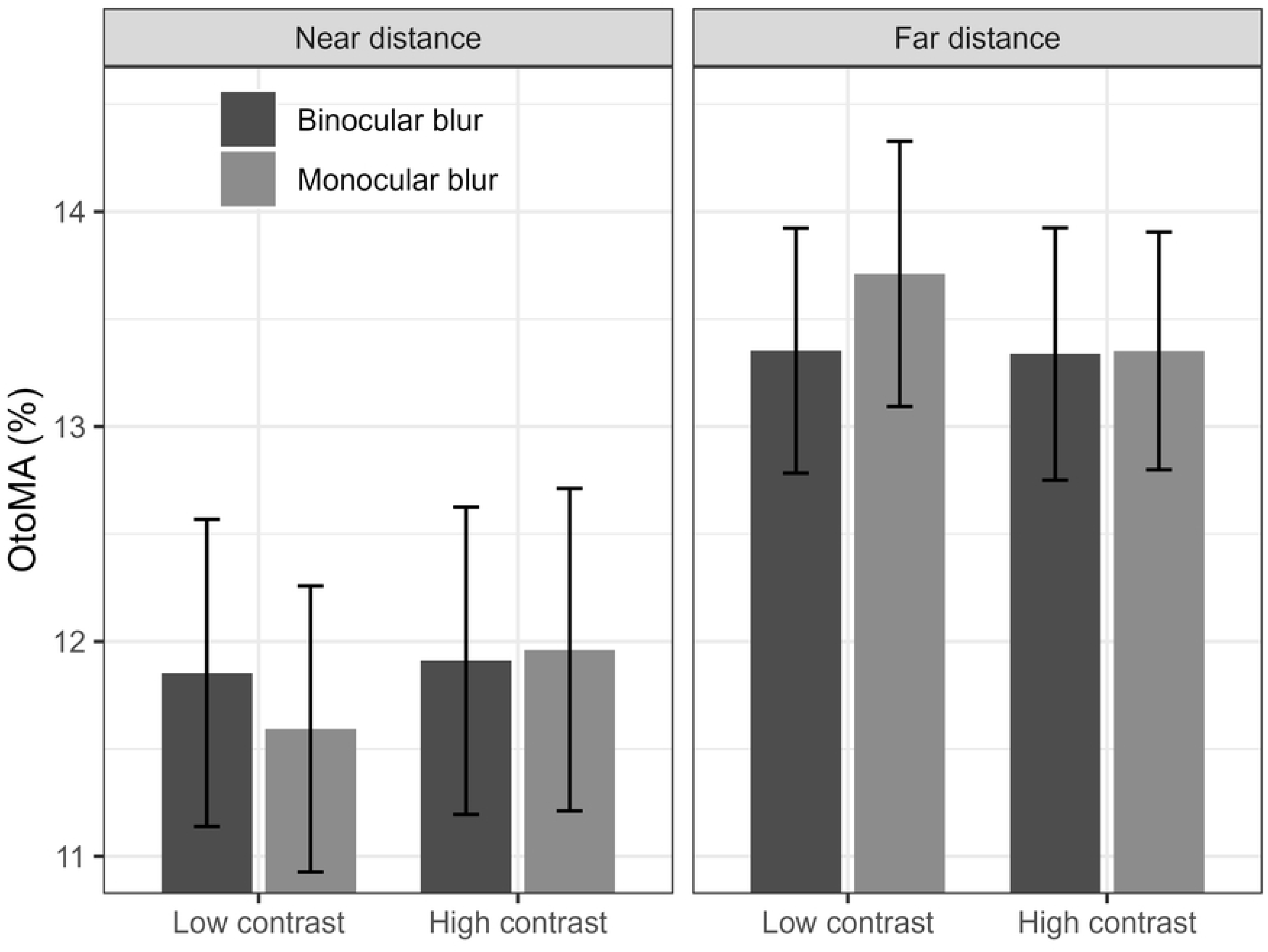
The effect of visual condition (binocular vs. monocular blur), contrast and distance (near vs. far) on the time from onset to maximum acceleration (% of total movement time). Error bars represent *SE*.

The baseline level of OtoMA showed substantial variability across participants, with the *SD* of the random intercepts estimated at 363.64% of the fixed intercept.

### Time from maximum acceleration to maximum velocity (MAtoMV)

A significant main effect of visual condition (monocular blur vs. full vision) indicated that full vision increased MAtoMV by 2.26% compared to monocular blur. The *SD* of the random intercepts was 131.82% of the fixed intercept, indicating high inter-individual variability in baseline levels of MAtoMV.

### Time from Onset to MGA (OtoMGA)

A significant three-way visual condition (binocular blur vs. monocular blur) x contrast x distance (near vs. far) indicated that monocular blur was associated with a slight increase in OtoMGA at near distance with high contrast targets and at far distance with low contrast targets, whereas, monocular blur was associated with a slight decrease in OtoMGA at near distance with low contrast targets and no apparent change in OtoMGA at far distance with a high contrast target (see Figure 8).

**Figure 8.**
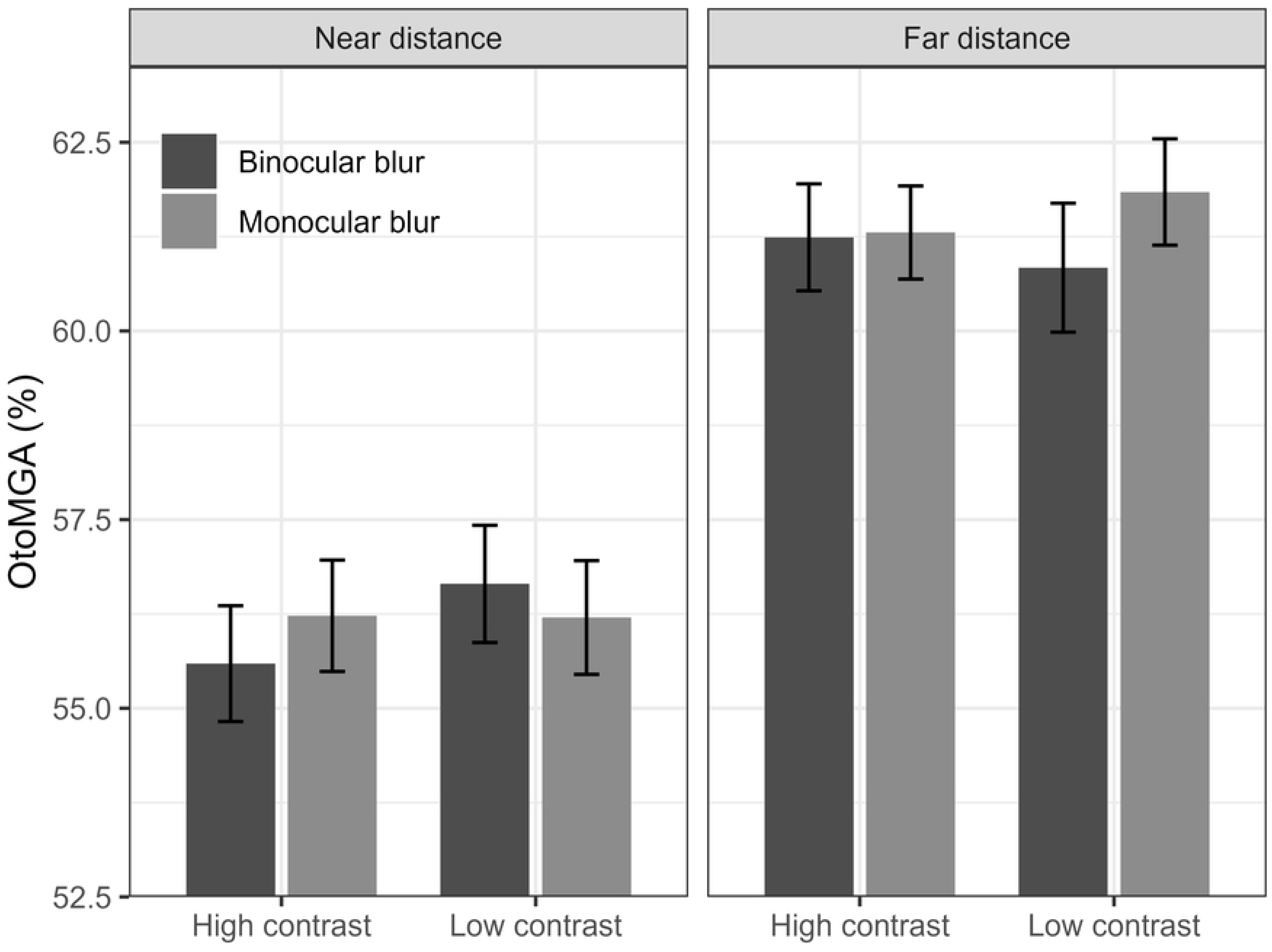
The effect of visual condition (binocular blur vs. monocular blur), contrast and distance (near vs. far distance) on the time to maximum grip aperture (% of total movement time). Error bars represent *SE*.

Similar to MGA, baseline levels of OtoMGA also showed high individual differences (*SD* of the random intercepts equal to 121.74% of the fixed intercept).

### Online control

The *β* values in Table 5 predict maximum deceleration, *µ_md_*, the time from maximum velocity to maximum deceleration, *µ_MV2MD_*, the time from maximum deceleration to contact with the target, *µ_MD2C_*, and dwell time, *µ_DwT_*. As the outcome variables are centred, the *β* values are expressed as a proportion of the measure’s overall *SD*, e.g., *µ_md_* is expressed as a proportion of the overall *SD* of MD (*SD_md_*).

### Maximum deceleration

No significant interactions or main effects were relevant to the study’s aims.

### Time from maximum velocity to maximum deceleration (MVtoMD)

A significant two-way visual condition (monocular blur vs. full vision) x distance (near vs. mid) interaction indicates that full vision is associated with an increase in MVtoMD at a near distance and a decrease at a mid distance (see Figure 9).

**Figure 9.**
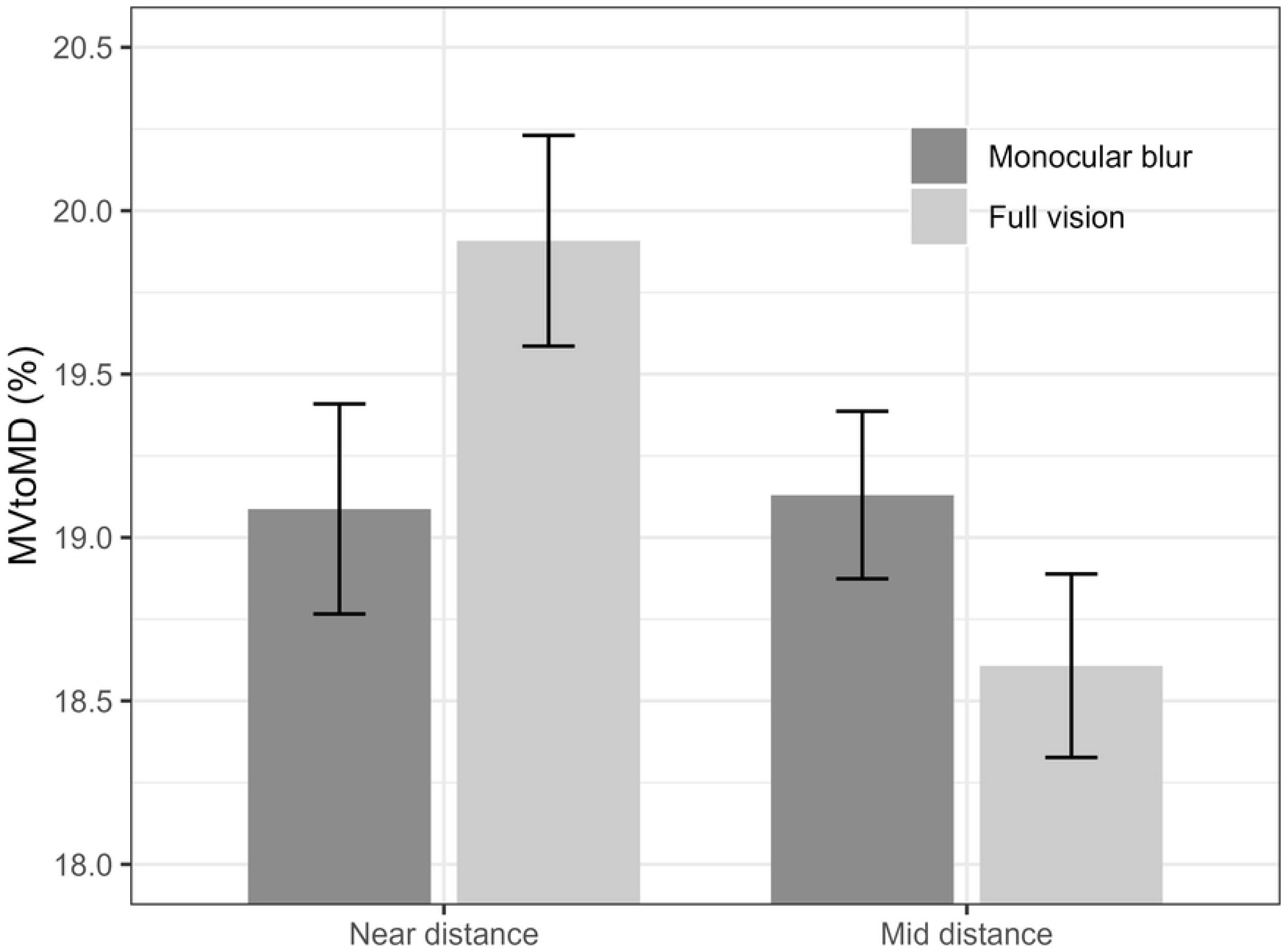
The effect of visual condition (monocular blur vs. full vision) x distance (near vs. mid) on the time from maximum velocity to maximum deceleration (% of total movement time). Error bars represent *SE*.

The *SD* of the random intercepts for MVtoMD showed high individual variability (110.34% of the fixed intercept).

### Time from maximum deceleration to contact with the target (MDtoC)

A significant two-way visual condition (monocular blur vs. full vision) x distance (near vs. mid distance) interaction indicated that full vision was associated with a decrease in MDtoC, relative to monocular blur, at a near distance and an increase in MDtoC at a mid distance (see Figure 10).

**Figure 10.**
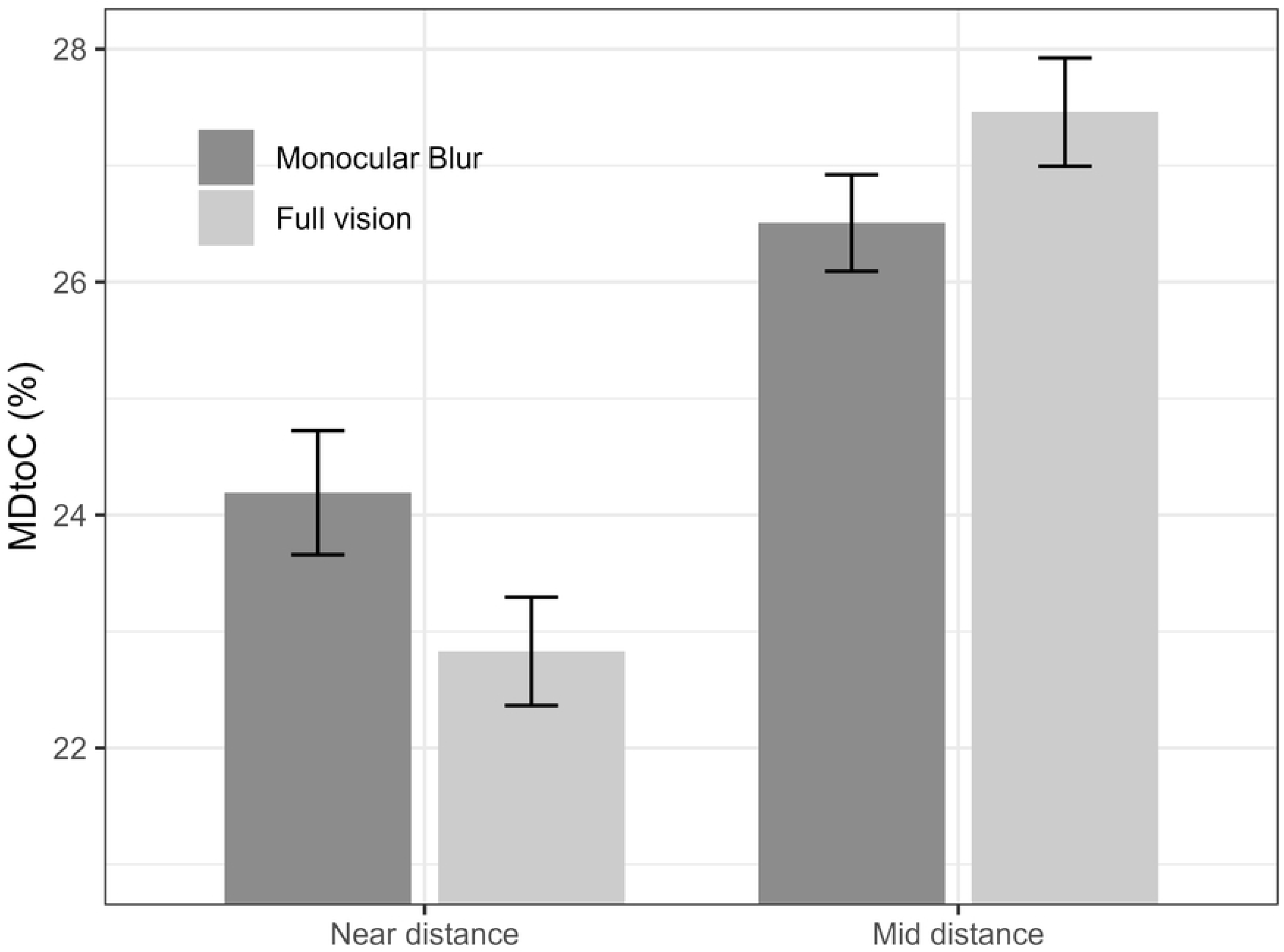
The effect of visual condition (monocular blur vs. full vision) and distance (near vs. mid) on the time from maximum deceleration to contact (% of total movement time). Error bars represent *SE*.

The *SD* of the random intercepts was 55.56% of the fixed intercept, indicating that the base level of MDtoC was moderately heterogeneous between participants.

### Time from contact to lift (dwell time [DwT])

MLM analysis revealed significant main effects of visual condition on DwT, which decreased progressively from binocular to monocular blur (5.08% lower) to full vision (3.68% lower) and by 12.54 mm/s^2^ and showed high individual differences in its baseline levels as indicated by the *SD* of the random intercepts being equal to 176.19% of the fixed intercept.

### Movement time

The *β* values in Table 6 predict movement time, *µ_mt_*. As movement time is centred, the *β* values are expressed as a proportion of movement time’s overall *SD*, *SD_mt_*.

**Table 6.** MLM estimates for fixed and random effects predicting online control characteristics of the RGL task.

| Estimate | Movement time |
| --- | --- |
| Intercept | 2.77***<br>[2.35, 3.19] |
| Three-way interactions ($\beta$) | Movement time |
| Binocular blur vs. monocular blur * Contrast * Near vs. mid distance | NS |
| Monocular blur vs. full vision * Contrast * Near vs. mid distance | NS |
| Binocular blur vs. monocular blur * Contrast * Near vs. far distance | NS |
| Monocular blur vs. full vision * Contrast * Near vs. far distance | -0.23*<br>[-0.45, -0.02] |
| Two-way interactions ($\beta$) | Movement time |
| Binocular blur vs. monocular blur * Contrast | NS |
| Monocular blur vs. full vision * Contrast | NS |
| Binocular blur vs. monocular blur * Near vs. mid distance | NS |
| Monocular blur vs. full vision * Near vs. mid distance | NS |
| Binocular blur vs. monocular blur * Near vs. far distance | NS |
| Monocular blur vs. full vision * Near vs. far distance | 0.16*<br>[0.01, 0.31] |
| Contrast * Near vs. mid distance | NS |
| Contrast * Near vs. far distance | NS |
| Main effects ($\beta$) | Movement time |
| Binocular blur vs. monocular blur | -0.27**<br>[-0.47, -0.06] |
| Monocular blur vs. full vision | -0.34***<br>[-0.49, -0.19] |
| Contrast | NS |
| Near vs. mid distance | 0.53***<br>[0.43, 0.63] |
| Near vs. far distance | 1.09***<br>[0.96, 1.22] |
| Stereoacuity | NS |
| CS | NS |
| VA | NS |
| Session number | NS |
| Random effects ($\sigma$) | Movement time |
| Participant ID | 0.21 |
| Near vs. mid distance | 0.09 |
| Near vs. far distance | 0.11 |
| Session number | 0.1 |
| Model fit | Movement time |
| R2 marginal | 0.736 |
| R2 conditional | 0.933 |

*Note:* Movement time and numeric predictors (VA, CS, stereoacuity, and session number) were standardised (i.e., with a mean equal to zero and *SD* equal to one). Movement time was modelled using a Gamma distribution. The Gamma distribution only accepts positive values; therefore, movement time was further transformed to be positive (movement time + 3). The numbers in each cell represent the estimate [lower 95% CI, upper 95% CI]. Blank cells (-) indicate that this predictor was not included in the final model. *NS* = $p \geq .05$ , \*\* = $p < .01$ , \*\*\* = $p < .001$ .

A significant three-way visual condition (monocular blur vs. full vision) x contrast x distance (near vs. far) indicated that full vision was associated with the largest decrease in MT at a far distance with low contrast targets (see Figure 11). There were also two main effects of visual condition: monocular blur reduces movement time by 32.80ms compared to binocular blur, and full vision reduces movement time by 41.93ms relative to monocular blur.

**Figure 11.**
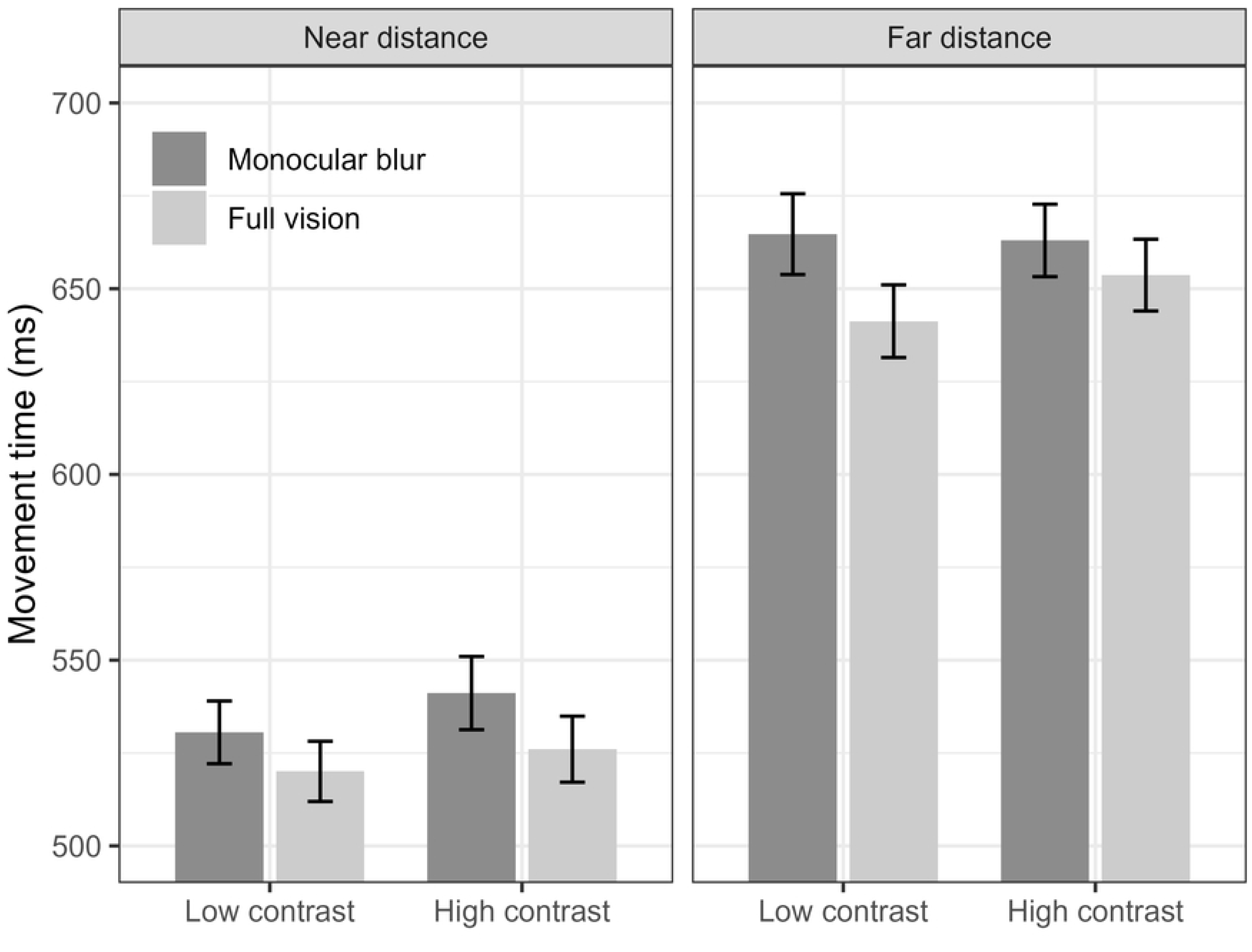
The effect of visual condition (monocular blur vs. full vision), contrast and distance (near vs. far) on movement time (s). Error bars represent *SE*.

The *SD* of the random intercepts was equal to 91.30% of the fixed intercept, indicating that the base level of MT was highly heterogeneous between participants.

## Discussion

The present study investigated the effects of monocular and binocular cataract-like visual blur and the interaction between visual blur and target contrast on the kinematic and timing characteristics of a reach-to-grasp task, marking a first in the literature. A priori, it was predicted that reducing visual blur would affect the scaling of the maximum grip aperture (MGA) and overall movement time (MT).

These effects would be coupled with increased deceleration time and dwell time (from target contact to lifting the target [DwT]). We also predicted that some, but not all, changes to performance associated with visual blur would be magnified by reducing the contrast of the targets. To match the progression of an individual undergoing cataract surgery, the effect of visual condition was estimated as the effect of monocular blur relative to binocular blur (as per first-eye cataract surgery) and the effect of full vision relative to monocular blur (as per second-eye cataract surgery).

To assess the effect of the visual condition on participants’ vision, contrast sensitivity (CS), stereoacuity, and visual acuity (VA) were tested under each condition. Modelling confirmed that monocular blur (compared to binocular blur) and full vision (compared to monocular blur) were associated with improvements in all three vision tests. This effect was largest for the transition from binocular blur to monocular blur, relative to the transition from monocular blur to full vision. The effect of visual condition on CS and VA was substantial (*R*^2^ marginal = 0.668 and 0.366, respectively); however, the effect of visual condition on stereoacuity was minimal (*R^2^ marginal* = 0.077; [Cohen, 1988]). At the time of writing, there appears to be no multilevel analysis of the visual outcomes of cataract surgery for comparison. However, these outcomes follow a similar pattern to those of previously published literature. For example, bilateral cataract removal was associated with the largest effect for CS (*r*^2^ = 0.46), followed by VA (*r*^2^ = 0.41) and the smallest effect for stereoactuity (*r*^2^ = 0.21; 44).

In line with research investigating the effects of amblyopia and monocular vision on prehension (1,6,10,11,14), superior vision (in this case, monocular blur relative to binocular blur, and full vision relative to monocular blur) was associated with a decrease in MT and with increased maximum acceleration (MA). No differences in maximum deceleration (MD) were found. This pattern replicates previous findings showing that control groups exhibit significantly greater MA but similar MD compared to amblyopia patients (14). Full vision relative to monocular blur did not affect maximum velocity (MV), while monocular blur relative to binocular blur increased MV at mid and far distances. A trend towards increased MV at a near distance did not reach significance, likely due to the lower speeds of smaller movements, in accordance with Fitts’ Law (45,46), where reduced movement magnitude limits the effect. These findings suggest that the improvement from binocular to monocular blur (analogous to first-eye cataract removal; FES) is sufficient to increase the MV in prehension movements, while the transition from monocular blur to full vision (second-eye cataract removal; SES) appears to provide no additional increase in MV.

Although the transition from monocular blur to full vision (SES) resulted in subtler improvements in prehension compared to the more dramatic effects of the transition from binocular to monocular blur (FES), gains in kinematic markers, such as reduced DwT and increased MA (which explain the reduction in movement time despite no change in MV), highlight that SES may be associated with critical benefits and enhancements in safety, efficiency, and independence during daily tasks.

The grip aperture (the distance between the thumb and index finger) is planned before movement initiation based on the object’s perceived size (47). Therefore, changes to grip apertures due to visual blur likely reflect changes in prehensile movements’ planning. In this study, monocular blur relative to binocular blur and full vision relative to monocular blur were associated with increases in MGA of 4.00mm and 3.16mm, respectively. These results contradict prior research showing that superior vision leads to no change in MGA in control participants compared to amblyopia patients (Grant et al., 2007) or with binocular vision compared with monocular vision (4,5,8). Similarly, the pattern contrasts with other studies reporting increased MGA with poorer vision, suggesting a greater safety margin (6,8,9). The increase in MGA associated with improved vision found in the present study may result from visual blur impairing the participants’ ability to accurately judge the target’s distance.

Servos et al. (1992) proposed that impaired vision (such as monocular vision compared to binocular vision) leads to participants underestimating target distance, which in turn causes the target to be perceived as smaller, resulting in smaller grip apertures. In the present study, the results suggest that improved vision was associated with increased estimates of target distance and, consequently, increased estimates of target size. This led to a larger MGA under monocular blur compared to binocular blur, and under full vision compared to monocular blur. This effect can be further evidenced by comparing MV with binocular and monocular blur. According to Fitts’s law, an increase in estimated distance should be associated with an increase in MV (45,46), which can be seen in Figure 5. Therefore, it seems reasonable to infer that participants estimated the target distance to be greater with monocular blur vs binocular blur, and scaled the target size accordingly at movement planning.

A point of note regarding the effect of visual condition on MGA is that, while the modelling suggests that monocular blur relative to binocular blur is associated with an increase in MGA, this is not represented in Figure 6, except for high contrast targets at a near distance. We considered the hypothesis that this effect might result from the multilevel structure of the analysis creating false positives or erroneous estimates. The effects of this multilevel structure were investigated by conducting a supplementary analysis (not reported in the results section). The simplest model of the effect of visual condition on MGA was calculated; it only included a main effect of visual condition and a random intercept for each participant. In this case, neither monocular blur relative to binocular blur nor full vision relative to monocular blur was associated with any change in MGA (*p* > .05).

However, when stereoacuity or VA was added to the model, the main effects of both monocular blur relative to binocular blur and full vision relative to monocular blur emerged, both associated with increases in MGA (*p* < .001, see https://github.com/willsheppard9895/blurNprehension/mgaModels.html for the model summary).

The effect of visual condition only emerged when individual differences in stereoacuity and VA were accounted for. This highlights two important considerations: first, that inter-individual differences in visual functions, such as VA and stereoacuity, evidenced by the analysis of random effects, may be masking the effects of visual impairment - in this case, visual blur. This demonstrates the need to consider each individual rather than applying a “one size fits all” methodology in eye care. Further support comes from the heterogeneity of performance on almost all motor outcomes in the RGL task, demonstrated by the analysis of the random intercepts. Second, this also suggests that other aspects of vision contribute to grip scaling beyond clinical measures. One possible explanation is vergence, which is crucial for grip aperture adjustments in prehension tasks (8). Unlike stereoacuity and VA, vergence behaviour is, at least in part, driven by visual blur, meaning that blur manipulation could impair vergence calibration, having a knock-on effect on depth perception (49) and potentially on visuomotor functions, as the observed changes in MGA strongly suggest.

Regarding the time course of prehensile movements, the present study showed that during the early/acceleration phase, monocular blur relative to binocular blur had no significant effect on the proportion of movement time spent reaching maximum acceleration (OtoMA) or transitioning from maximum acceleration to maximum velocity (MAtoMV). Similarly, full vision relative to monocular blur was also associated with no overall change in OtoMA. However, we found a significant 2.26% increase in MAtoMV, suggesting that SES might be associated with improved planning of prehensile movements. The increase in the percentage of overall movement time in the early stages of the movement (OtoMA and MAtoMV) suggests that full vision reduces the number of corrections being made in the later stages of the movement. These results contrast with studies that found no effect or an absolute decrease in acceleration time associated with superior vision (see 14 for an example). On the other hand, the increase in MAtoMV observed here is comparable in scale to previous findings when expressed as a percentage of total movement time, 1.62% (9).

In the late/deceleration phase, there were no main effects of visual condition on either the time between max velocity and max deceleration (MVtoMD) or from MD to first contact with the target (MDtoC). The impact of visual condition interacted with near versus mid distance, whereby full vision relative to monocular blur was associated with an increase in MVtoMD at a near distance and a decrease at mid distance (see Figure 9), compared with a decrease in MDtoC at a near distance and an increase in MDtoC at mid distance (see Figure 10). However, these effects are small (∼1% difference) and not coupled with a main effect, so they seem inconsequential practically. As reducing visual blur has a limited effect on the time course of the late/deceleration phase of the movement, this suggests that reducing visual blur is associated with lesser changes to the online control of prehensile movements (50) compared to the significant impact of reducing visual blur on metrics associated with movement planning (such as MA, MV and MGA).

Although there was no main effect of visual condition on the time from onset to MGA (OtoMGA), a significant three-way interaction suggested that full vision (relative to monocular blur) increased OtoMGA under the most challenging conditions (far distance, low contrast). However, considering the variability in previous findings from the literature (−7.80% to +5.11% change in OtoMGA with binocular vs. monocular vision) and the overall positive effects of full vision on movement planning (e.g., increased MA, MV, and reduced dwell time), this isolated interaction is unlikely to indicate a meaningful impairment in planning.

As per previous research, monocular blur and full vision (relative to binocular and monocular blur, respectively) were associated with decreased dwell time (9,10,14). A decrease in dwell time suggests that improving vision improves the representation of the object’s position and physical parameters, reducing the reliance on somatosensory feedback from the finger and thumb and allowing the lift to be executed more quickly, analogous to reduced online control errors (10). This result is particularly compelling when coupled with the increases in MA (monocular blur and full vision), MV (full vision only), and MGA (monocular blur and full vision). It seems reasonable to conclude that reducing visual blur leads to better planning and online control of prehensile movements. Reducing visual blur allows individuals to plan better prehensile movements, most likely due to changes in the perception of the shape/size of the target. This is coupled with an improved ability to make in-flight adjustments, facilitating the efficient and safe performance of the movement.

Future studies could benefit from using liquid crystal glasses to control the precise timing of visual input. Grant and Conway (2019) demonstrated that removing visual feedback after movement initiation removed any binocular advantage associated with full vision compared to amblyopia; it would be beneficial to know if this also applies to visual blur. This, in turn, would help us better understand the situations in which patients with conditions such as cataracts may face the greatest difficulties with prehension. Furthermore, it would be good to manipulate visual cues to delineate effects outside of CS, VA and stereoacuity (the effect of visual blur on stereoacuity was very small, *R*^2^ = 0.077, see Table 3). For example, Loftus et al. (2004) manipulated the availability of vergence information, allowing us to test the theory suggested earlier in the discussion. Furthermore, future research may want to manipulate blur levels to gain an estimate of the point at which visual blur and visual conditions such as cataracts may begin to make prehensile tasks increasingly difficult.

A critical limitation of this study is that our setup did not effectively control for monocular depth cues: without a chinrest, participants might compensate for visual blur by adjusting their head position and viewpoint (although this is not entirely atypical, see 9). This ability to adapt is directly relevant to real-world cataract patients, particularly those who have undergone FES and retain one unimpaired eye. FES patients can partially mitigate the functional limitations of monocular blur through compensatory head movements and behavioural adjustments, reducing their reliance on degraded stereo and vergence cues. However, this compensatory strategy does not replicate the experience of true binocular clarity. The results of this study reinforce that even when individuals have access to monocular adaptation strategies, movement planning and control continue to improve with full binocular vision. SES provides functional advantages that extend beyond behavioural compensation, reinforcing its clinical importance.

The results proposed here, along with previously published work, suggest that reducing visual blur and, by proxy, timely FES and SES improve the planning and, to a lesser extent, the online control of prehensile movements. These findings motivate a future programme of work to investigate the effects of FES and SES in cataract patients as they move through their surgery journey, thus providing a better understanding of the potential benefits of cataract surgery on skilled action.

## Acknowledgements

Thanks to everyone that helped.

